# Infant gut microbiomes contribute to metabolic states that impact brain function

**DOI:** 10.64898/2026.03.09.710596

**Authors:** Firas S. Midani, Do-Hun Lee, Younghye Moon, Maggie Seale, Thomas D. Horvath, A. Kyle Ardis, José Cantú, Emavieve Coles, Jason D. Pizzini, Duolong Zhu, Sean W. Dooling, Grace J. Ahern, Colleen K. Ardis, Alisha Beckford, Nicole M. Ruggiero, John Shin, Raphaela Joos, Catherine Stanton, R. Paul Ross, Darlene L.Y. Dai, Piushkumar J. Mandhane, Charisse Petersen, Stuart E. Turvey, Mairead E. Kiely, Deirdre M. Murray, Mauro Costa-Mattioli, Kimberley F. Tolias, Robert A. Britton, Heather A. Danhof

**Author notes:** Senior author (Robert A. Britton, Heather A. Danhof). Correspondence (F.S.M), (R.A.B), (H.A.D).

## Abstract

Alterations in the gut microbiome are associated with neurodevelopmental disorders, but causal mechanisms and therapeutic strategies remain undefined. Here, we demonstrate that human infant microbiomes isolated during the first six months of life drive behavioral impairments in mice and that microbiota-based interventions restore mice to normal behavior. Early-life microbiomes from twelve infants who later exhibited cognitive deficits at 2 years old (low-scoring) transferred adverse metabolic, brain, and behavioral phenotypes to mice, in contrast to microbiomes from twenty-three cognitively typical or high-scoring infants. Deficits in mice were rescued by fecal microbiota transplant from high-scoring infants or a rationally designed consortium that promoted amino acid levels. We confirmed lower fecal amino acid concentrations in low-scoring infants and replicated the association between early-life microbiome composition and cognitive outcomes in a second geographically independent infant cohort. Altogether, we discovered an early-life microbiome-mediated metabolic state causally linked to cognitive deficits and amenable to microbial intervention.

## INTRODUCTION

Over the past two decades, the re-discovery of the human microbiome revealed several important connections between the bacteria living in/on the body and human disease. One area of intense investigation has been the impact of the gut-brain axis on brain function, which uncovered connections of the microbiome to numerous adult-onset neurological disorders^1–4^. The ability of microbes to alleviate brain disorders has also been shown in mice; for example, *Akkermansia muciniphila* and *Parabacteroides distasonis* promote seizure protection in epilepsy through alterations of gamma-glutamylated amino acids, glutamate, and gamma-aminobutyric acid (GABA)^5^. Despite extensive evidence linking the microbiome to neurological disorders, the consequences of bacterial variation during infancy, a critical period of brain development, on neurodevelopmental delays and disorders (NDDs) remains elusive and controversial.

Nearly 1 in 6 children is diagnosed with an NDD, including autism spectrum disorder, intellectual disability, and attention-deficit/hyperactivity disorder^6^. Individuals with NDDs exhibit alterations in their fecal microbiome as infants^7,8^. Large longitudinal studies of children, followed from birth and extending into early/late childhood, identified associations between specific microbial taxa and/or metabolites with neurodevelopmental profiles. For example, increased levels of the *Bacteroides* genus were associated with higher cognitive performance in multiple studies^9–11^ and were shown to ameliorate autism-related abnormalities in mice^12^. The microbe *Limosilactobacillus reuteri* was shown to alleviate sociability deficits in multiple animal models of autism, across multiple laboratories^13,14^ and improved sociability in children diagnosed with autism^15,16^ demonstrating the therapeutic potential of microbiota-based interventions for NDDs. These studies highlight an exciting opportunity to understand if the infant gut microbiome might play a causal role in shaping brain development and a potential to design microbial-based interventions for children at risk for NDDs.

Despite many groundbreaking discoveries in the microbiome field, there has been little progress in defining whether and how human gut microbes modulate CNS-driven behaviors. To address this gap, we therefore pursued an unprecedented approach designed to overcome long-standing limitations of microbiome studies^17,18^. Our study began with a human infant cohort study, progressed through extensive mechanistic investigations in murine models, and culminated in replication of findings in a second human infant cohort. Humanized microbiota mouse (HMM) models, in which human fecal samples are transferred into germ-free mice, remain the gold-standard for revealing causality between human microbes and impact on human health. However, most HMM studies use low numbers of human fecal samples and lack longitudinal sampling across subjects. As such, studies often use mice as the biological units, rather than the subjects’ fecal samples, which should be treated as the true biological units in these studies^18^. This practice has contributed to an implausible level of success reported for HMM studies and overstated the role and causality of the microbiome in human disease. To address this, we increased the number of human fecal samples in our HMM study, rather than solely depending on large numbers of mice to identify significant phenotypes. We also sought to replicate key findings linking the early-life gut microbiome to NDDs in a second independent human birth cohort.

To determine the impact of the early-life microbiome on behavioral outcomes, thirty-five infants were stratified based on a cognitive development test administered at 2 years old and categorized into either low-scoring, typical-scoring, or high-scoring. We transferred 115 longitudinally collected infant fecal samples between 1 and 12 months of age into germ-free mice to generate HMM lines, one for each fecal sample, then studied behavior and analyzed brains in the resulting offspring from each line. We observed that every fecal sample collected from low-scoring infants during the first 4 months of life caused multiple behavioral and cognitive deficits in animals. We were able to reverse these phenotypes by introducing microbiomes from high-scoring infants into low-scoring animals. Multi-omics revealed metabolites that are reproducibly reduced in mice generated from low-scoring infants. Consequently, we used metabolic modeling to rationally design a simple, defined consortium of bacteria that corrected the metabolic deficiency and behavioral problems. To demonstrate clinical relevance, we confirmed metabolic signatures in the infant samples, and, in a second infant cohort from a different continent, we replicated the association between gut microbiome structure and neurodevelopmental outcomes at 2 years old. These results demonstrated that infants who experience cognitive and behavioral deficits later in life may have early-life microbiomes that contribute to their developmental delay.

## RESULTS

### Infant gut microbial composition fails to predict cognitive outcomes at two years of age

To investigate whether the early-life gut microbiome influences neurocognitive development, we analyzed data from the Cork Nutrition and Development Maternal-Infant Cohort (COMBINE), which longitudinally tracked 456 Irish healthy term born infants from the third trimester of pregnancy to 2 years old. Infant fecal samples were collected at six timepoints throughout the first year, and comprehensive clinical metadata were collected on mothers and infants after birth and during follow-up visits (Figure 1A). At the age of 2 years, 297 infants completed cognitive testing using the Bayley Scales of Infant and Toddler Development, Third Edition (BSID-III), yielding a normally distributed composite cognitive score (104.1 ± 14.5, Figure 1B). For a subset of 129 infants, we performed whole-genome shotgun sequencing on 435 longitudinally collected fecal samples. As expected in early life, we detected an increase in microbial alpha diversity over time and beta diversity was associated with age and other covariates including delivery method, gestational age, and feeding behavior (Figure S1A–B). Interestingly, cognitive BSID-III scores at 24 months were negatively correlated with alpha-diversity at 4 and 6 months and positively correlated at 12 months (ρ = −0.37, −0.39, and 0.32, Figure S1C–D) and likewise beta diversity correlated with BSID-III scores at 1, 4, and 6 months (R^2^ of 0.029, 0.031, and 0.027, Figure S1E). We compared the abundance of microbial species and pathways with BSID-III scores but did not detect any statistically significant correlations, although we did detect species and pathways that are significantly associated with infant age and other covariates (Table S1). Because infant age may confound associations of cognitive scores with the microbiota, we also performed simpler analyses that compared microbiota within narrower temporal windows, 1–4 months or 9–12 months (Table S1), but still did not detect any significant associations of microbial species or pathways with BSID-III scores.

**Figure 1.**
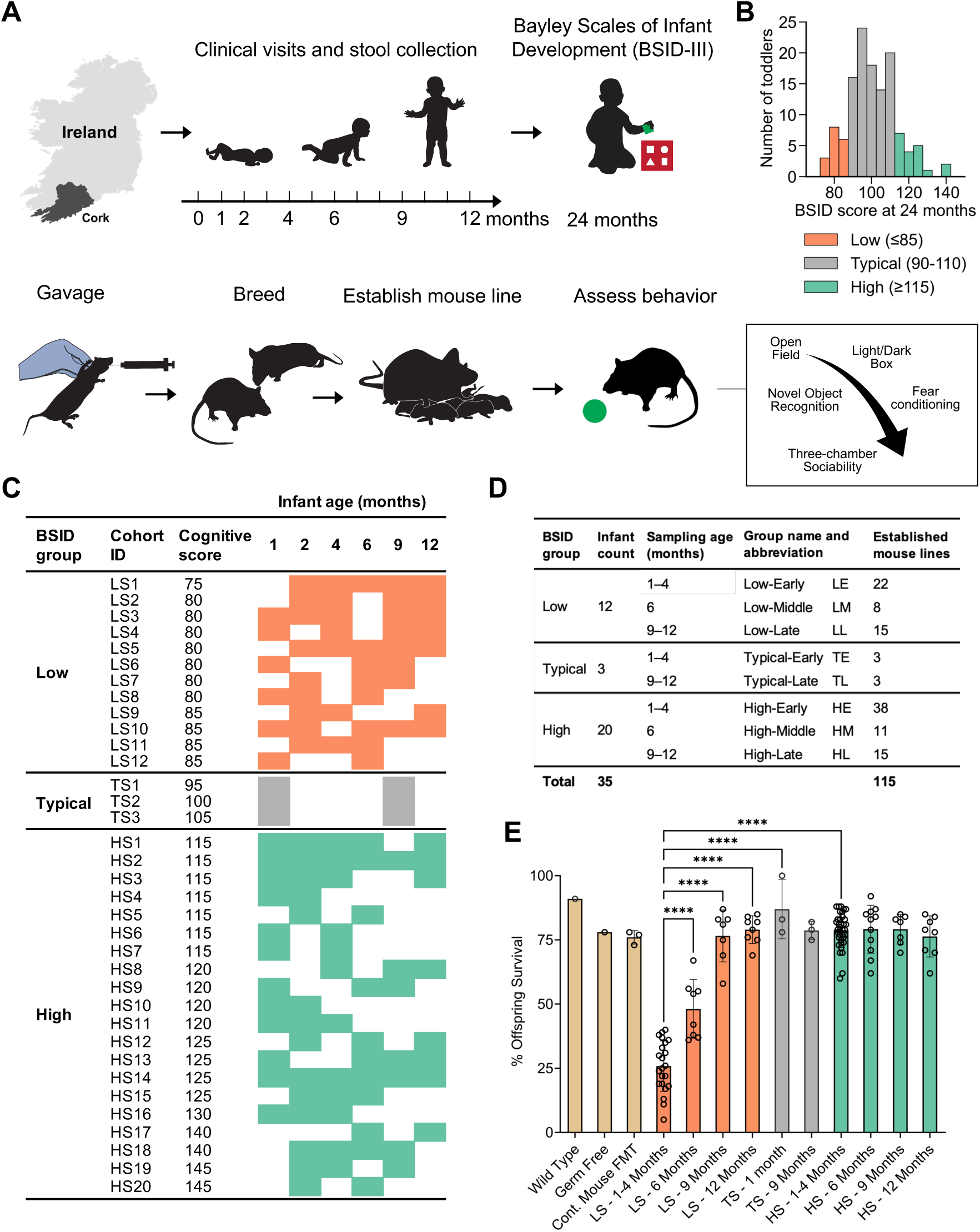
Study design and lower postnatal survival in mice humanized with microbiota from low-scoring infants. (A) Stool samples were collected from infants in the Irish COMBINE study. Humanized-microbiota mouse colonies were generated from each infant fecal sample by gavaging the sample into pairs of germ-free mice, which were subsequently bred to maintain mouse lines and generate offspring for behavioral assessments. (B) Infants were stratified into low, typical, or high-scoring groups based on BSID-III cognitive scores. (C) Table of infant samples used to establish humanized microbiota mice stratified by age at time of collection and BSID-III scores. (D) Established mouse colonies were stratified into groups based on the donor’s age at time of stool collection and BSID-III scores. (E) LE mice displayed significantly lower postnatal survival compared to other groups. Tan=controls, orange=low-scoring (LS), gray=typical-scoring (TS), green=high-scoring (HS).****p < 0.0001 by one-way ANOVA with n=102 mouse lines.

Given the limited interpretability of BSID-III scores within its typical range^19^, we next focused our analysis on infants with the highest and lowest cognitive scores. We identified a subset of 35 infants with BSID-III cognitive scores and at least two fecal samples collected during the first year of life (Figures 1C–D). This final cohort included 12 low (BSID-III ≤ 85), 3 typical (BSID-III 95–105), and 20 high (BSID-III ≥ 115) scoring individuals. Low-scoring infants had higher Caesarean delivery rates and infant formula use (p < 0.05), while sex, gestational age, and household income were comparable between low and high-scoring infants (Table S2). To generate humanized microbiota mouse (HMM) lines from these infants (n=115 samples), we created fecal slurries, re-sequenced them, and repeated our microbial analyses comparing low and high-scoring infants at the strain level. Consistent with our broader cohort, we detected statistically significant associations of BSID-III scores with alpha-diversity at 1 and 6 months (Figure S2), but strains and pathway abundances were not predictive of BSID-III scores (Table S3). Although we detected modest correlations of alpha- and beta-diversity with BSID-III scores, these analyses suggest that broad taxonomic and metagenomic profiles alone do not robustly predict cognitive performance at two years old in this cohort.

### Early-life microbiomes from 1–4 months-old low-scoring infants impaired rearing behavior

To assess the impact of the infant microbiome on behavior and cognition, we created stable mouse lines for each of the longitudinally collected fecal samples from 35 infants. In total, 115 mouse lines were established by orally gavaging each fecal sample (FMT) into 8-week-old germ-free mating pairs, including 45 low-scoring (LS), 6 typical-scoring (TS), and 64 high-scoring (HS) infant fecal samples (Figures 1C–D). HMM lines were later grouped by infant age at sample collection, with those from 1-4 months termed “early”, those from 6 months as “middle”, and those from 9-12 months as “late”. As controls, we compared these lines to conventional wild-type mice, germ-free mice, and germ-free mice that received an FMT from a wild-type mouse.

Interestingly, during the establishment of gnotobiotic mouse lines, multiple fecal samples yielded breeding pairs that struggled to rear pups to weaning. Observational monitoring of breeding pairs humanized with samples from low-scoring infants revealed increased anxiety-like, inattentive, and/or aggressive behaviors relative to other groups. Notably, LE mice frequently failed to engage in normal nesting behavior or display maternal care, and pups often did not survive beyond early postnatal stages.

Therefore, we systematically tracked litter sizes and pup survival in all 115 breeding lines. There were no differences in litter sizes amongst breeding pairs of different groups (Figure S3A). However, all mice receiving FMT from LS infant samples collected at 1–4 months of age had < 50% pup survival over multiple litters and generations, despite having similarly-sized litters (Figure 1E). Notably, no difference in pup survival occurred in mice given an FMT collected between 9-12 months from these same infants, which was similar to other groups. Lastly, a transitional survival rate was observed for animals given 6-month LS FMT compared to 6-month HS FMT. As subsequent data demonstrated that mice colonized with LS infant samples had consistent results over all 12 infants, we will refer to 1–4 months fecal samples as Low-Early (LE), 6 months as Low-Middle (LM), and 9–12 months fecal samples as Low-Late (LL). For comparison, typical- and high-scoring infant samples will be similarly referred to as TE, TL, HE, HM, and HL. These findings indicate that the infant microbiome of LE lines robustly and reproducibly transferred adverse behavioral phenotypes to mice, in contrast to other groups, and point to a critical window during which the infant gut microbiome impacts behavior.

### LE microbiota induced anxiety-like behavior

To further define the adverse phenotypes transferred by the gut microbiome, we performed an extensive assessment of mouse behaviors using four well-established tests: open field, light/dark box, novel object recognition, and fear conditioning. For these assessments, a total of 98 mouse lines were tested, and each line was assessed with groups of 16–24 mice (half male and half female) from generations F1–F3. A subset of mouse lines with 12 mice each (half male and half female) were also tested for sociability and social novelty.

To investigate whether early-life microbiota influences emotional reactivity, we assessed anxiety-like behavior using the open field test. This test evaluates locomotor activity and anxiety by monitoring mouse movement in a square open area during a 10-minute period. Total locomotion is quantitated to assess the gross motor deficits that would confound interpretation of many behavioral tests. All groups displayed comparable levels of movement. However, during the same test, time spent away in the center from the box walls is used as a measurement of anxiety. LE mice spent significantly less time in the center of the arena relative to all other groups, indicating increased anxiety (Figure S3B). Like pup rearing outcomes, LM mice spent significantly more time than LE lines in the center but significantly less time than HM mice (Figure S3B), further highlighting that the first 6 months of life is a critical window for gut microbiome development.

The light/dark box test was performed to further assess anxiety in the animals. In this test, half of the compartment is enclosed (“dark”) and half is open (“light”). Mice are curious and will typically explore the entire area, however, reduction of total test time spent in the light side is an indication of anxiety. Consistent with the open field test, LE mice also displayed increased aversion to the illuminated compartment compared to control or other humanized-microbiota lines (Figure 2A, S3C). Altogether, these findings indicated that LE microbiota promotes an anxiety-prone state in mice.

**Figure 2.**
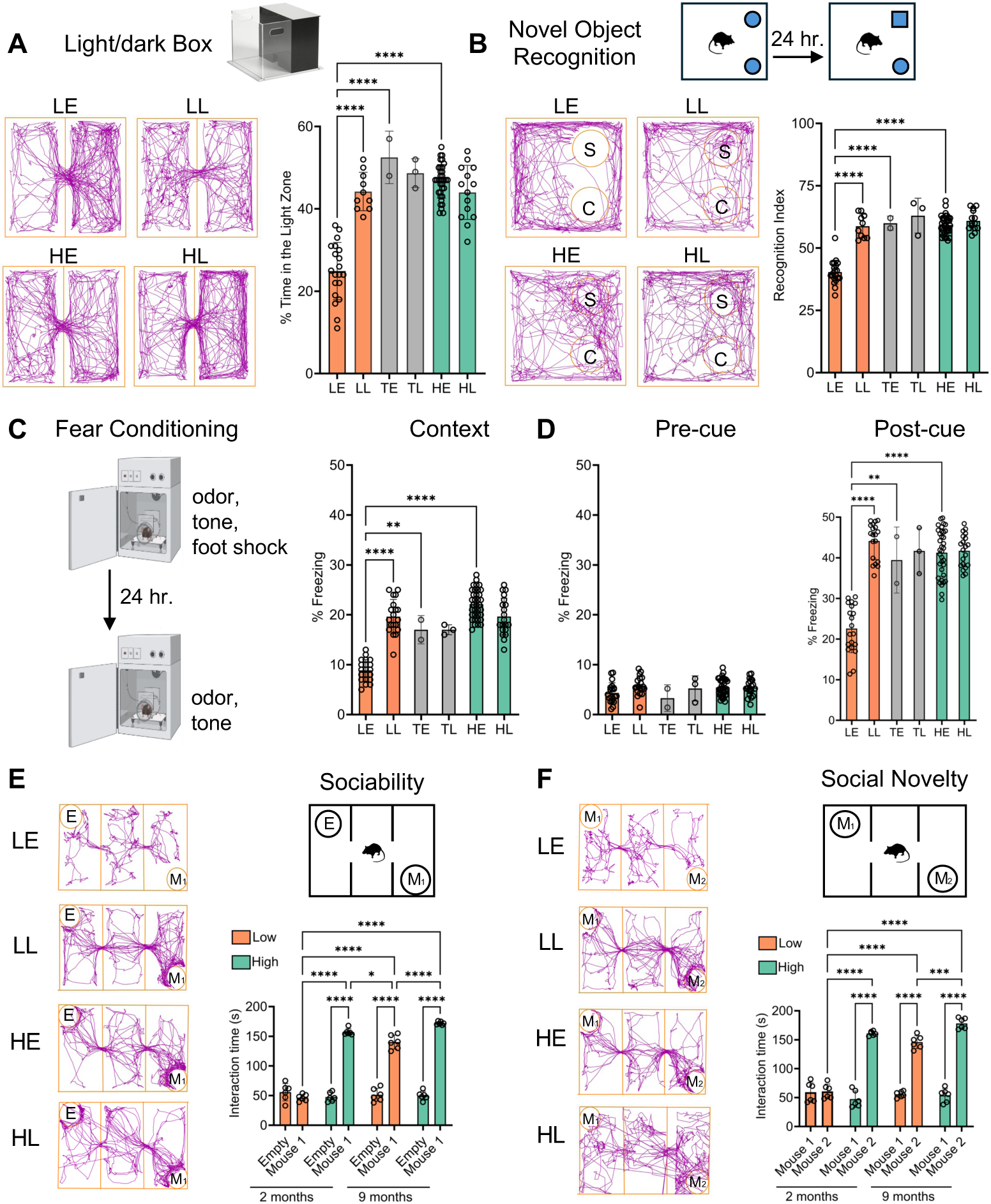
LE mice display anxiety-like behavior and impairments in learning, memory, and social functions. (A) Percentage of total test time of 10 minutes spent in the light compartment of the light/dark box behavioral test. Representative traces of the movement of individual mice are shown. Each circle in the graph represents the average score of a mouse line. (B) Percentage of total test time of 10 minutes spent with the novel object in the novel object recognition test. Time spent with the novel object/total time was calculated and multiplied by 100 to determine the recognition index. Representative traces of the movement of individual mice are shown. Each circle represents the average score of a mouse line. (C) Percentage of total test time spent freezing during 5 minutes of contextual fear conditioning, which is the proxy for memory of an adverse event environment. Each circle represents the average score of a mouse line. (D) Percentage of total test time immobile during the first 120 seconds (pre-cued) prior to tone and 120 seconds after tone (post-cued) of fear conditioning, which is the proxy for memory of adverse events following an auditory tone. No differences were observed between groups in pre-cued. (E) Total time spent interacting either with an empty cup or with a mouse during a sociability test. Each point is an individual mouse; n=4 groups. (F) Total time spent interacting with either with a familiar mouse (mouse 1) or a stranger mouse (mouse 2) during a sociability test. Each point is an individual mouse; n=4 groups Behavioral icons (A and C) are sourced from ANY-maze and BioRender.com. Orange=low-scoring, gray=typical-scoring, green=high-scoring. In panels A–D, 1 point=24 mice, n=98 mouse lines. In panels E–F, n=6 mice/group tested, n=4 groups. Data are presented as a mean ± SD. *p < 0.05, **p < 0.01, ***p < 0.001 **** p≤ 0.0001 by one-way ANOVA.

### LE mice exhibit learning and memory deficits

Next, we assessed cognitive performance using novel object recognition (NOR) and fear conditioning. The NOR task assesses recognition memory in mice by evaluating their innate preference for novelty. Mice were familiarized with two identical objects in an open field, followed by a test phase the next day where one familiar object was replaced with a novel one. Interaction with each object is quantitated, and higher exploration with the novel object indicates intact recognition memory. All groups except LE mice demonstrated the expected preference for the novel object, as indicated by recognition indices well above 50%. In contrast, LE mice failed to show novelty preference and instead spent more time investigating the familiar object (Figures 2B, S3D). This reversal of preference in LE mice may reflect an impaired recognition memory and/or heightened anxiety-related avoidance of unfamiliar stimuli.

Given the deficits in object recognition memory, we next assessed associative fear memory using a standard Pavlovian fear conditioning paradigm. In this task, mice were placed in a conditioning chamber and exposed to a neutral stimulus (e.g., environment and auditory tone) that ended with an aversive stimulus (e.g., mild foot shock). Freezing behavior is used as a proxy for associative memory. We found that LE mice displayed significantly reduced freezing in the training chamber 24 hours after training compared to all other mouse groups (Figures 2C, S3E), indicating an impaired recall of the training environment.

After the mice have been measured for contextual memory and have resumed movement, the tone is played again and amount of time the animals freeze is measured. LE mice again displayed significantly lower freezing in response to the tone than all other mouse groups (Figures 2D, S3F), indicating a defect in cued memory. Importantly, no differences were observed in pre-tone baseline freezing across groups, confirming intact sensorimotor function and further supporting the conclusion that LE microbiota impairs both hippocampal-dependent contextual memory and hippocampal-independent cued memory.

### LE mice exhibit impaired social behaviors

Because social dysfunction is a core feature of several neurodevelopmental disorders, we assessed sociability and social novelty preference using the three-chamber social interaction tests. Here, mice are placed in a chamber that contains two cups, one holds another mouse and the other is empty. Mice will typically spend more time interacting with the cup containing another mouse versus the empty cup. During the sociability phase, all mouse groups except LE mice displayed a robust preference for interacting with the mouse versus the empty enclosure, as would normally be expected. In contrast, LE mice showed no preference, spending equal time with the mouse and the empty cup (Figure 2E), indicating that sociability is impaired in these mice. We then assessed social novelty, in which a novel mouse is introduced into the cup opposite to the now-familiar mouse. Typically, mice will spend more time with the novel mouse than the familiar mouse. We found that all non-LE mouse groups preferentially interacted with the novel mouse, while LE mice failed to discriminate between the novel and familiar mice (Figure 2F). These findings demonstrate that LE microbiota disrupted both social approach and social novelty recognition, resembling phenotypes observed in mouse models of autism spectrum disorder and other conditions marked by early-life social impairments.

### LE mice exhibit disrupted synaptic architecture and widespread neuroinflammation

To further investigate how early-life gut microbiota impacts brain development, we performed immunohistochemical analyses on the mouse brains following behavioral testing. Guided by behavioral evidence of hippocampal and cortical dysfunction, we examined synaptic organization in the dentate gyrus (DG) and cornu ammonis 1 (CA1), two hippocampal regions essential for learning, memory, and mood regulation, and the somatosensory cortex, which processes sensory information from the body^20–23^. Inhibitory synaptic development was assessed using the postsynaptic marker Gephyrin. LE mice exhibited significantly elevated Gephyrin expression in both the DG and CA1 compared to HE mice, consistent with increased inhibitory synaptic drive within hippocampal circuits (Figures 3A–B). In contrast, the excitatory postsynaptic marker PSD95 did not differ significantly between LE and HE mice in either hippocampal region (Figures 3C–D). No differences in Gephyrin or PSD95 were observed in the somatosensory cortex (Figures S4A–B), and presynaptic inhibitory vesicular GABA transporter (VGAT) and excitatory vesicular glutamate transporter (VGLUT) markers were comparable across all three brain regions (Fig. S4D–I), suggesting that these synaptic alterations are region-specific and predominantly postsynaptic. Because proper hippocampal function relies on a tightly calibrated balance of excitatory and inhibitory (E/I) signaling, this LE-associated shift towards increased inhibition is notable and aligns with E/I disruptions frequently observed in neurodevelopmental disorders^24,25^. To assess neuromodulatory tone, we also examined serotonin (5-HT), a neuromodulator important for mood regulation, cognitive flexibility, and memory formation^26–28^. Immunostaining revealed a marked reduction in 5-HT levels in both the hippocampus and somatosensory cortex of LE mice (Figures 3E–F, S4C), consistent with their elevated anxiety-like behaviors and memory deficits. These findings indicate that microbiota from low-scoring infants disrupted multiple aspects of synaptic organization, including E/I balance and serotonergic input, which likely alter hippocampal circuit dynamics.

**Figure 3.**
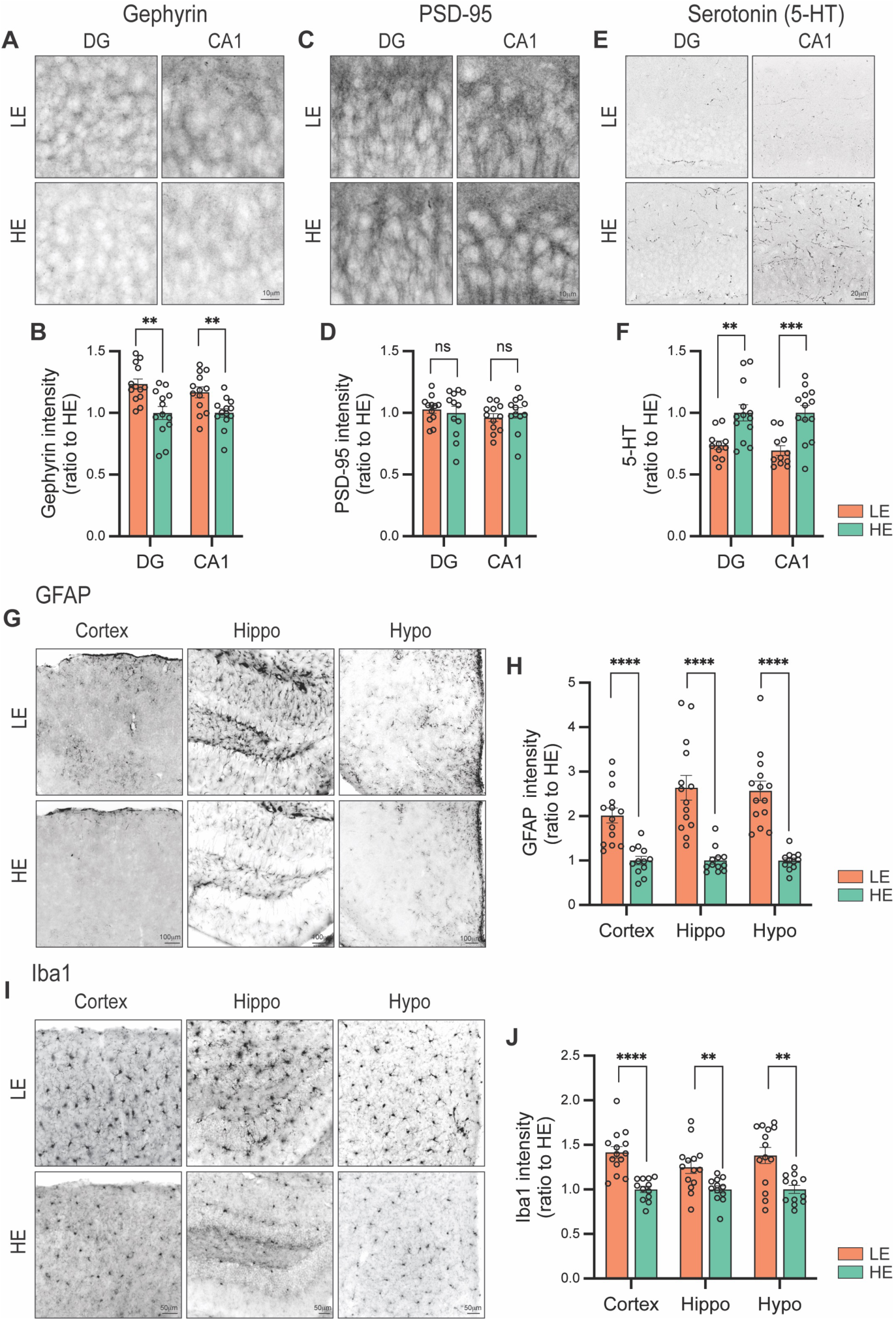
Brain immunohistochemistry and imaging reveal aberrant synaptic architecture and widespread neuroinflammation in LE mice. (A–B) Inhibitory synaptic development assessed in the dentate gyrus (DG) and CA1 hippocampal regions using the postsynaptic marker Gephyrin showed increased Gephyrin staining in LE mice relative to HE mice. (C–D) Excitatory synaptic development evaluated using the postsynaptic marker PSD95 in hippocampal regions DG and CA1 showed no difference between LE and HE mice. (E–F) Serotonin (5-hydroxytryptamine, 5-HT) immunoreactivity in hippocampal DG and CA1 regions. LE mice exhibited a marked reduction in 5-HT staining intensity in both regions relative to HE mice. (G–H) Astrocytic activation assessed by GFAP staining intensity in the cortex, hippocampus (Hippo) and hypothalamus (Hypo). GFAP levels were increased across all brain regions in LE compared to HE mice. (I–J) Microglial activation based on Iba1 staining intensity in the cortex, hippocampus, and hypothalamus. Iba1 intensity was markedly elevated in all examined brain regions in LE mice relative to HE mice. Data are normalized to the HE group and presented as a mean ± SEM. Ns, not significant, **p < 0.01, ***p < 0.001, ****p < 0.0001 by unpaired Student’s t-test.

Because astrocytes and microglia are essential regulators of synapse formation, function, and circuit refinement, and prolonged glial activation can impair these processes^29–31^, we next assessed neuroinflammatory status. Using glial fibrillary acidic protein (GFAP) as a marker for reactive astrocytes, we detected widespread astrogliosis in the hippocampus, cortex, and hypothalamus of LE mice, whereas GFAP levels remained low in HE brains (Figures 3G–H). Microglial activation, assessed via ionized calcium-binding adapter molecule 1 (Iba1) staining, was likewise robustly increased across all three brain regions in LE mice relative to HE mice (Figures 3I–J). This neuroinflammatory profile reflects a pathological state in which glia cells adopt chronically reactive phenotypes marked by morphological activation, heightened phagocytic activity, and increased release of inflammatory mediators capable of disrupting synapse maturation and circuit refinement^32–34^. Together, these results suggest that microbiota from low-scoring infants elicited persistent neuroinflammation alongside synaptic disorganization, two features strongly associated with adverse neurodevelopmental outcomes. These convergent alterations in inhibitory tone, serotonergic signaling, and glial reactivity suggest a potential link between early-life microbiome and the behavioral abnormalities observed in LE mice.

### Gut microbial composition in mice does not predict donor cognitive outcomes

Given the consistent behavioral and brain differences between LE mice and other groups, we sought to identify microbial features associated with behavioral and brain phenotypes in the mice. Our first approach was to profile their gut microbiomes with whole-genome shotgun sequencing, and test for differences in microbial diversity, microbial strain composition, and pathway abundance. Unlike the microbiomes of their infant donors, there was no difference in alpha-diversity or beta-diversity between LE and TE/HE mice (Figure S5A), and none of the strains or pathways were significantly different between these groups, although we detected differences associated with the infant donor age (Figure S5B; Table S4). Despite our low statistical power, we also tested for differential abundance of microbial genes which were quantified and classified by short-read profiling into KEGG orthogroups, EC enzyme classes, and Pfam protein families. Again, we detected significant associations of microbial genes/functions with donor age but not donor cognitive outcomes (Table S4). Finally, we tested whether ten neuroactive pathways are significantly enriched or depleted in LE microbial metagenomes using feature set enrichment analysis of KEGG orthogroups. We only identified significant depletion in LE mice for genes involved in the synthesis of tryptophan, an essential precursor for the neurotransmitter serotonin (Figure 4A; Table S4). In summary, we did not detect robust or broad differences between LE and HE mice either because our analysis was underpowered or because the adverse impacts of LE microbiota are driven by collective microbial metabolism and interspecies interactions that are not captured at the species, gene, or pathway levels. Hence, gut microbial composition alone is insufficient to predict donor cognitive outcomes.

**Figure 4.**
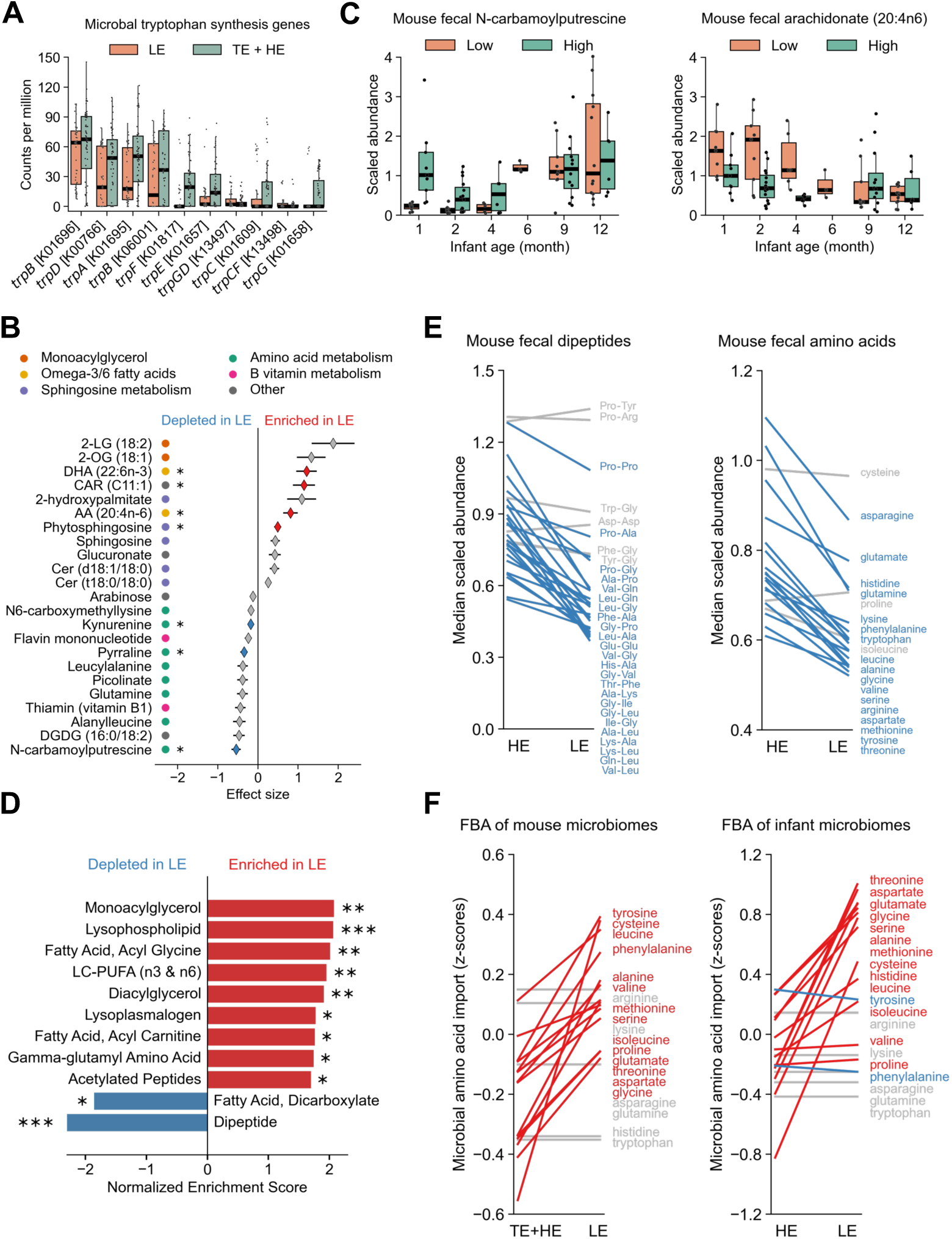
Microbiome-driven metabolic state depletes amino acids in LE mice. (A) Relative abundance of microbial tryptophan synthesis. (B) Fecal metabolites that were differentially altered in LE mice (p < 0.01). Colored markers group metabolites based on chemical relatedness or similarity. (C) Abundances of two representative metabolites, one enriched and the other depleted in LE compared to LL mice yet both are comparable between LL and HL mice. (D) Metabolic groups that were significantly depleted or enriched in LE mice. (E) Median abundance of all dipeptides and amino acids detected in mouse feces with untargeted metabolomics, comparing HE and LE mice only. (F) Median z-scores for microbial reactions that import amino acids in flux balance analysis (FBA) models. 2-LG=2-linoleoyl glycerol, 2-OG=2-oleoyl glycerol, AA=arachidonic acid, CAR=carnitine, Cer=ceramide, DGDG=digalactosyldiacylglycerol, DHA=docosahexaenoic acid, LC-PUFA=long-chain polyunsaturated fatty acid. Data is presented as either median ± interquartile range (A and C) or mean ± SEM (B). In panels A and C, 1 point=1 mouse. Blue and red lines indicate metabolites with median abundances or z-scores at least 10% lower or higher, respectively, in LE compared to HE subjects. *p < 0.05, **p < 0.01, ***p < 0.001 by linear mixed effects models (B) or feature set enrichment analysis (D) and adjusted with the Benjamini-Hochberg method (B and D).

### Gut microbial metabolism in mice is associated with donor cognitive outcomes

Because differences between LE and HE mice may reflect altered microbial metabolic activity not captured by species- or gene-based analysis, we next performed untargeted metabolomics on mouse feces. Strikingly, we identified 7 significantly altered metabolites, including ones classified as sphingosines, amino acids, and long-chain polyunsaturated fatty acids (LC-PUFAs) (Figures 4B–C, S5D; Table S5). Based on feature set enrichment analysis, where metabolites are grouped based on chemical similarity, LE mice also displayed significant depletions of dipeptides and dicarboxylates and enrichments of gamma-glutamyl amino acids, acetylated peptides, and a variety of fatty acids including monoacylglycerols, LC-PUFAs, acyl glycines, acyl carnitines, lysophospholipids, and lysoplamalogen (Figure 4D; Table S5). Remarkably, 24 out of 30 dipeptides detected in mouse feces had decreased abundances in LE relative to HE mice, and 17 out of 20 amino acids displayed a similar pattern (Figure 4E). These results demonstrated a major metabolic shift in the fecal contents of LE mice.

To ascertain if the mouse gut microbiomes contribute to these metabolic differences, we compared the metabolic potential of LE and TE/HE microbiomes using several genome-scale methods. First, we pivoted our microbial metagenomic approach from profiling short reads to assembling genomes in mouse fecal samples. Then, we mined these metagenomes for metabolic gene clusters, which comprised physically clustered genes involved in the production of primary metabolites central for cell survival and growth, and detected consistent depletion for primary metabolic pathways in LE mice, including pathways related to amino acids, peptides, fatty acids, fumarate, pyruvate, and acetate metabolism (Figure S5E; Table S6). Second, we modeled microbial metabolism for each community with flux balance analysis using publicly available genome-scale metabolic reconstructions. In concordance with untargeted metabolomics, these microbial community-scale metabolic models simulated higher rates for microbial consumption of amino acids in LE mice, with higher rates for 14 out of 20 amino acids in LE mouse guts (Figure 4F). Remarkably, metabolic modeling of the infant microbiomes also simulated higher consumption of 13 amino acids in LE infant guts (Figure 4F). Additionally, models estimated lower cross feeding in LE mice for leucine, serine, acetate, and hydrogen ions (Figure S5F; Table S6), the latter suggesting higher acidification in the lumen of LE mice. Altogether, these results demonstrated that LE gut microbiomes encode fewer primary metabolic pathways and are more likely to deplete amino acids in the gut which could explain the major metabolic shift observed in LE mice. Importantly, amino acid availability is tightly coupled to the synthesis of key neurotransmitters and may therefore be related to the disrupted synaptic architecture and neuroinflammation observed in LE brains.

### Fecal microbial transplants from high-scoring donors restore normal phenotypes in LE mice

We next asked if the adverse behavioral and metabolic phenotypes in LE mice can be reversed by high-scoring infant microbiomes. We selected three LE mouse lines as FMT recipients and three HL infant samples as FMT donors (Figure 5A). In each combination, 3-week-old LE mice received an HL FMT via a single oral gavage. We then compared the behavior of these mice to corresponding parental lines as well as autologous transplant control lines, which are LE backgrounds that received a second FMT of the same donor microbiome. We observed that all HL samples rescued breeding, and behavior deficits of all LE recipient lines tested (Figures 5B, S6A). In contrast, all three FMT autologous control lines retained their adverse breeding and behavioral phenotypes in the light/dark box (Figures 5C, S6B) and fear conditioning tests (Figures 5D, S6C–D).

**Figure 5.**
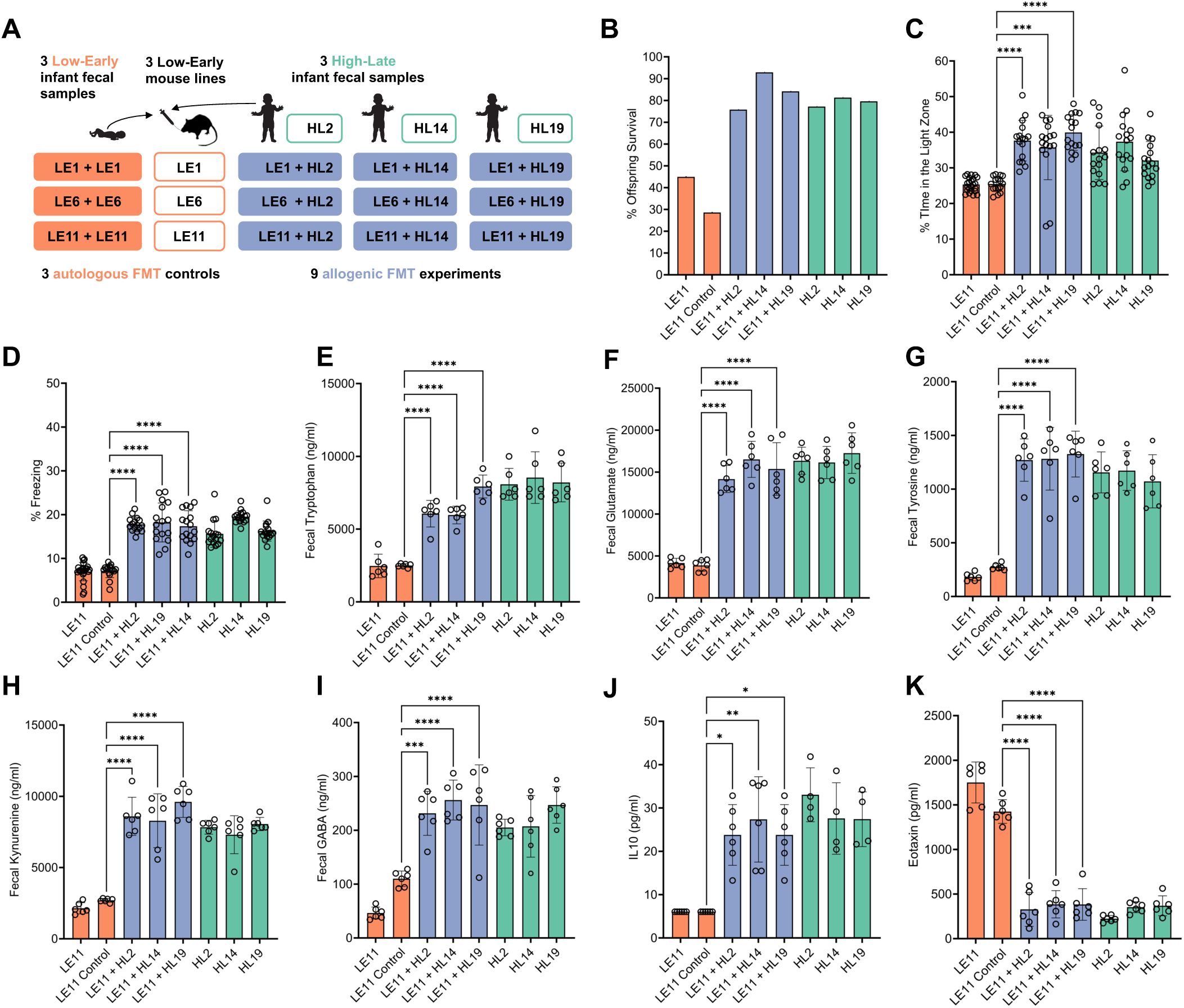
Metabolic, immune, and behavioral phenotypes are restored by FMT of high-scoring infant stool into LE mice. (A) Experimental design for transfer of a second fecal slurry into humanized mice to determine if microbiota could ameliorate phenotypes. (B) Pup survival returned to similar levels as HL animals when LE mice were given a second FMT at weaning. (C) Mice underwent the light/dark box behavioral test, for a total of 10 minutes and the time in the light compartment was quantitated. Allogenic FMT mice spent more time exploring the light side than LE controls. (D) Mice that received a second. Allogenic FMT mice froze more frequently in the contextual fear test than LE controls. (E–I) Fecal levels of tryptophan, glutamate, tyrosine, kynurenine, and GABA increased to levels comparable to HL animals. (J–K) Serum levels of proinflammatory eotaxin decreased in LE animals given a second FMT from HL infants. Levels of IL-10 increased after second FMT to similar levels of HL mice. IL-10 was not detectable in LE mice. Orange=low-scoring, blue=mice given a second FMT from HL infants, green=high-scoring. 1 point=1 mouse, n=16–24 mice per group. Data are presented as mean ± SD. *p < 0.05, **p < 0.01, ***p < 0.001, **** p≤ 0.0001 by one-way ANOVA.

Moreover, we observed that FMTs can also reverse adverse metabolic and immune markers in LE mice. Using targeted metabolomics, we observed that successful FMT rescues were marked by increased abundance of amino acids, including tryptophan, glutamate, and tyrosine (Figures 5E–G, S7A-C). In contrast, amino acid concentrations did not change in autologous self-rescue controls. Further, neuroactive metabolites derived from amino acids, including kynurenine and GABA (Figures 5FH-I, S7D–E), also remained unchanged in controls but increased significantly in successful rescues.

To assess systemic inflammation, we profiled 32 immune markers in mouse plasma. LE mice showed reduced levels of the anti-inflammatory cytokine IL-10, and increased levels of proinflammatory eotaxin (CCL11), both of which were restored to HL parental levels by HL FMT transfer (Figures 5J-K, S7F-G). In control lines, both eotaxin and IL-10 levels remained unchanged. Overall, these results demonstrate that LE mice have altered behavioral, metabolic, and immune phenotypes that can be rescued by microbial transplants from high-scoring infants.

### Rationally designed microbial consortium restores normal phenotypes in LE mice

Based on the multi-omic analyses in our animal study, we proposed a mechanistic model for a microbiome-driven metabolic state that impairs behavior. We hypothesized that LE microbiomes significantly deplete amino acids from the intestinal environment, thereby reducing amino acid uptake by the host and disrupting immune and brain pathways that rely on amino acid-derived metabolites and neurotransmitters. To identify infant-derived bacteria that promote neurotypical brain function, we focused on species that are most correlated with the metabolic environment associated with HE microbiomes and species capable of synthesizing amino acids or promoting their levels in the gut. First, we correlated the top 7 metabolites from our differential analysis of untargeted metabolomics of mouse feces with all species in the corresponding humanized mouse microbiomes. We then ranked species based on their number of significant (p < 0.05) correlations with HE-enriched metabolites and anticorrelations with LE-enriched metabolites (Figures 6A, S8A). This data-driven approach identified several species associated with the favorable metabolic state in high-scoring microbiomes. Second, to identify species capable of synthesizing most or all amino acids, we annotated biosynthetic pathways for 17 amino acids in microbial genomes assembled from mouse samples (Figure 6B). Finally, we revisited our community-scale metabolic modeling and identified species that most robustly reduced consumption of amino acids and promoted cross feeding of key metabolites (Figures 6C, S8B–C). Altogether, we used genomic evidence, microbial community-scale simulations, and microbial-metabolite correlations to identify and rank species that may restore normal metabolic states.

**Figure 6.**
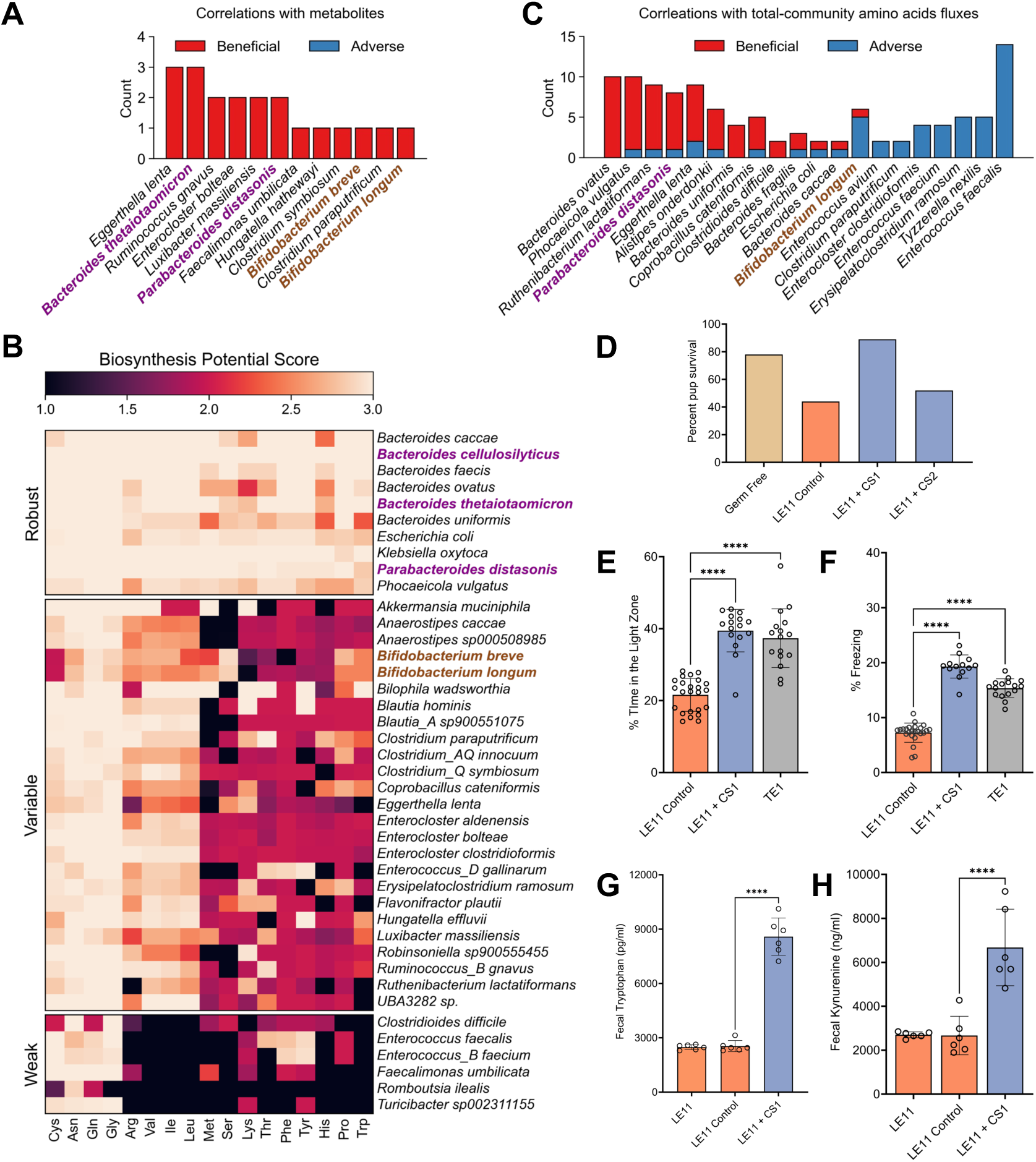
Rescue of adverse metabolic and behavioral phenotypes with a rationally-designed microbial consortium. (A) Number of significant correlations between microbial strains and seven metabolites that are significantly enriched or depleted in LE mice. Correlations were tested using median abundances within each mouse colony. (B) Scores for amino acid biosynthetic potential by the most prevalent microbial species in mice. Scores of 1, 2, and 3 correspond to low, medium, and high confidence for encoding a complete pathway for the synthesis of an amino acid. (C) Number of significant correlations between microbial species and community-scale amino acids fluxes in flux balance models. (D) Pup survival increased significantly when animals were given CS1 but not CS2 twice weekly. (E–F) Twice weekly gavage with CS1 restored behavioral measures to levels comparable to TE mice in the light dark box (E) and contextual fear conditioning (F) tests. (G–H) Levels of fecal metabolites tryptophan (G) and kynurenine (H) increased to normal ranges following twice weekly treatment with CS1. Orange=low-scoring, blue=mice given a second FMT from HL infants, green=high-scoring. Species labels in purple and brown indicate members of CS1 and CS2 respectively. In panels A–C, analysis was based on n=28 mouse lines; n=3–4 mice per line (A), 1,747 genomes (B), and 207 metabolic models (C), and correlations were computed using Spearman’s test (A) or linear mixed effects models (C) and denoted as significant if p < 0.05. In panels E-H, 1point=1 mouse, n=16–24 mice per group. Data are presented as a mean ± SD (E–H). In panels E-F, *p < 0.05, **p < 0.01, ***p < 0.001, **** p≤ 0.0001 by one-way ANOVA.

Finally, we aimed to reverse metabolic, breeding, and behavioral phenotypes in LE mice using a simple microbial consortium rationally designed by our data-driven approach. First, we isolated 120 microbial strains (73 species) from high-scoring infant fecal samples, then created a three-species microbial consortium (referred to as CS1) comprised of species that scored highly in our ranking assays*: Bacteroides thetaiotaomicron*, *Bacteroides cellulosilyticus,* and *Parabacteroides distasonis*. As a control, we designed a second consortium (CS2), using species commonly employed as infant probiotics, which our modeling indicated would be unable to robustly improve amino acid concentration (Figures 5A–C, S8A–C). CS2 was also comprised of three strains isolated from high-scoring infants: *Bifidobacterium breve*, *Bifidobacterium longum* subspecies *suillum*, and *Bifidobacterium longum* subspecies *infantis*. CS1 or CS2 was orally gavaged into 3-week-old LE mice. As expected, CS2 did not improve breeding success. In contrast, CS1 significantly increased post-natal pup survival, improved behavioral outcomes (increased time in the light zone in the light/dark box test and higher freezing rate in the fear conditioning tests), and elevated amino acid and kynurenine concentrations, (Figures 6D–H). In summary, a rationally designed three-member microbial consortium reversed metabolic, breeding, and behavioral phenotypes in LE mice.

### Infant samples from both Ireland and Canada confirm microbial and metabolic signatures

To establish the human relevance of our preclinical findings, we compared infant samples from two independent cohorts in Ireland and Canada. These human datasets provided critical validation of the microbial and metabolic pathways identified in our mouse models. First, we compared microbial diversity in infants from this study’s Irish COMBINE cohort^35^ with those from the Canadian CHILD cohort^36^. Across both populations, early-life (3–4 month) alpha diversity was negatively correlated with BSID-III cognitive scores at 24 months (Figure 7A), demonstrating a robust and reproducible human microbial signature. Second, we measured fecal amino acid concentrations in the Irish early-life infant samples. Low-scoring infants had significantly depleted levels of glutamine and GABA, closely mirroring the metabolic patterns observed in infant-humanized mice (Figure 7B). This concordance provides direct human support for the metabolic state inferred from our animal studies. Logistic regression trained on glutamine concentrations also robustly discriminated LE and HE fecal samples, with an area under the curve of 97.9% and an operating point corresponding to a sensitivity of 90.4% at a specificity of 99% (**Figure 7C**). Together, these results demonstrate alignment between our preclinical findings and clinical evidence, which strongly supports amino acids as early diagnostic biomarkers for infants at higher risk of neurodevelopmental delays.

**Figure 7.**
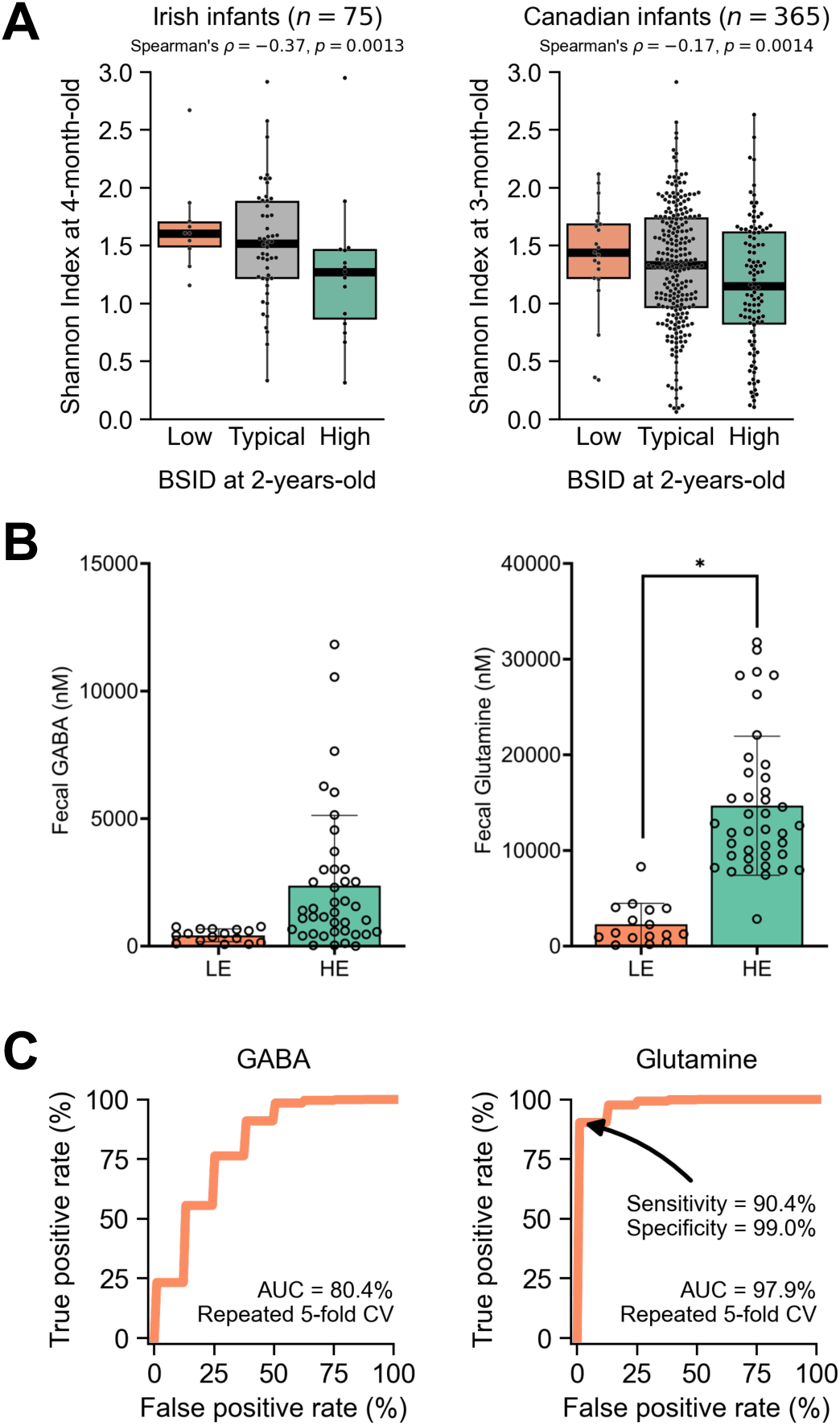
Microbial diversity signature is validated in an independent Canadian cohort and metabolic biomarkers discriminated Irish infant donors by their cognitive scores. (A) Microbial Shannon index in the COMBINE and CHILD study cohorts in Cork, Ireland and Edmonton, Canada, respectively, stratified by infant cognitive scores on the BSID-III test at 2 years old. (B) Concentrations of GABA and glutamine in infant stool samples from the Irish COMBINE cohort. Each point is an individual fecal sample (LE n=16, HE n=40). (C) ROC curves for predictive models using logistic regression on 100 repeats of stratified 5-fold cross-validation, displaying the average area under the curve (AUC) across all cross-validation tests. Data are presented as a median ± interquartile ranges (A) or mean ± SD (B). Statistical significance was determined by Spearman’s correlation on the continuous BSID-III scores (A) or one-way ANOVA (B). (*p < 0.05).

## DISCUSSION

Neurodevelopmental disorders and/or delay impact about 1 in 6 children in the United States^6^. Although NDDs have been associated with genetics, environment, diet, and the gut microbiome, underlying causes remain poorly understood. Using an unprecedented approach, we showed that fecal microbiomes from infants in the first 4 months of life, have profound effects on brain function and behavior when transplanted into germ-free mice. The consistency of the results was remarkable, with every fecal sample from LE infants (22 independent biological samples) repeatedly displaying similar adverse impacts on behavior and cognitive functions. Based on these behavioral assessments in mice, we can identify which infants had a BSID-III cognitive score at least 1 standard deviation below the mean at two years of age. These results demonstrate that fecal microbial communities associated with low cognitive ability in infants in the first 4 months of life also influence brain function in mice.

Although many clinical surveys have associated the infant gut microbiome with cognition and neurodevelopment, causal demonstrations have been lacking until now. While these surveys identified taxonomic associations of microbes with neurodevelopmental delay^7,9,37^, we did not detect any significant species associations, likely due to the small sample size of our cohort and the high inter-individual variation of microbiome composition. Instead, we identified functional features in the microbiome that drive metabolic differences between LE and TE/HE mice, including the striking reduction of amino acids and dipeptides in LE animals. An independent humanized microbiota study, which sampled infants and stratified them by cognitive scores at 6 months old, observed a similar trend of decreased histidine in low-scoring animals^10^. In addition to supporting core biosynthetic, metabolic, and epigenetic processes, amino acids serve as neurotransmitters and precursors of neurotransmitters and neurodevelopmental signaling molecules, including GABA and kynurenine that were identified in this study. Notably, alterations in glutamate (excitatory neurotransmitter), GABA (inhibitory neurotransmitter), and tryptophan (serotonin precursor) levels detected in our metabolomic analyses are consistent with the changes we observed in synaptic markers in the brain, pointing to a potential mechanism of action. In an early life stress mouse model, microbially-mediated reduction of glutamine in plasma indeed impacted glutamine levels in the brain and induced social dysfunction and anxiety-like behaviors, which were reversible by *L. reuteri* treatment^38^. Other possible mechanisms, that are not mutually exclusive, may play a role, because we identified additional metabolic and immune markers impacted by the LE microbiome. Notably, eotaxin, which was increased in LE mice, has been implicated in neurological defects associated with long Covid and directly impairs brain function when administered systemically to mice^39,40^. Microbial supplementation, either with a whole-community FMT or a simple defined consortium, reversed behavioral phenotypes and corrected both metabolic and immune differences, so further investigation is needed to define the potentially overlapping mechanisms of action. Altogether, we propose that early-life infant microbiomes associated with low cognitive scores have a common metabolic state: a preference to consume amino acids when compared to microbiomes that are associated with typical/high-scoring infants. Our metabolic modeling supports this hypothesis, and a rationally-designed consortium of bacteria was able to restore key metabolites and completely reverse behavioral deficits. While we found a similar microbial signature of alpha diversity in an independent Canadian cohort, additional validation of features and biomarkers are needed to determine the breadth of this microbiome-driven metabolic state.

Interventions for NDDs are often limited to managing symptoms after diagnosis, with developmental surveillance focused on infants with genetic risks, prematurity, significant neonatal illness or brain injury. Approximately 15-20% of children will display low test scores on neurodevelopmental tests such as BSID-III at 2 years old, and 15% of children in low- and middle-income countries (LMICs) fail to meet cognitive milestones at 3-4 years old ^41^. The BSID-III is a developmental assessment, giving a single cognitive score, and serves as a surrogate marker for later intellectual ability with stronger prediction in high-risk cohorts. In low-risk cohorts such as COMBINE, the BSID-III has a low sensitivity, however, the specificity of a low BSID-III score for later cognitive difficulties is high; >94% for an IQ score 1 standard deviation below the mean at 5 years of age. Our results suggest that the infant microbiome and the proposed metabolic state may serve as an early biomarker of later cognitive outcomes, enabling identification of children at elevated risk for neurodevelopmental disorders in the first 6 months of life, thereby offering opportunities for early intervention. One promising candidate for clinical diagnosis of the metabolic state is glutamine, which was depleted in fecal samples from low-scoring infants as young as 1-month-old. Despite these samples being in cold storage for several years, glutamine levels in infant stool were predictive of high versus low BSID-III scores with 90% sensitivity and 99% specificity. Ultimately, our findings establish a powerful framework for the early identification of at-risk infants and the deployment of microbial interventions designed to promote healthy brain development and prevent NDDs during a critical developmental window.

### Limitations of the study

We observed impaired behavior in animals colonized with microbes from infants with low BSID-III scores, but it is unclear whether these behavioral observations would be considered a neurodevelopmental phenotype in the animals. These phenotypes also coincided with an altered metabolic state, but more work is needed to determine if the altered metabolites are a mechanistic link between the gut and impaired brain function. Although we showed clear metabolite alterations in infant fecal material, we could not measure circulating levels in the infant donors because we lacked blood samples. Future studies are also needed to determine if levels of these metabolites are altered in the brain and how they impact brain phenotypes that we observe. This study demonstrated that these metabolic alterations are driven by gut microbiomes with a preference for consuming amino acids, but it is unknown which environmental factors select for these microbiomes. While we propose microbial interventions to correct for the adverse metabolic state, a complementary approach is needed to identify and address the maternal, prenatal, and postnatal factors that predispose the development of these adverse microbiomes. Future birth cohorts could apply this complementary approach and simultaneously validate the microbiome-driven metabolic state in other countries and cultures, especially because the majority of our work was based on a single infant cohort in a Western developed country. More importantly, our approach identified children who score poorly on BSID-III tests at 2 years old, but a more selective approach that identifies infants who are later diagnosed with NDDs will be more clinically relevant. Long term outcomes beyond 2 years was not available to this study, but this would be important to examine in later studies, with outcomes at least at school age. Finally, our work identified a critical age window, from birth to 6 months of age, where the gut microbiome impacts brain development, but it is unknown if intervention is possible after this time to correct for microbial impact on behavior. Altogether, our study discovered a microbiome-driven metabolic state associated with cognitive outcomes, but additional research is needed to validate this metabolic state, its etiology, prognosis, and mechanism of action in terms of neurodevelopment and brain function.

## RESOURCE AVAILABILITY

### Lead contact

Further information and requests for resources and reagents should be directed to and will be fulfilled by the lead contact, Robert A. Britton.

### Materials availability

This study did not generate new unique reagents.

### Data and code availability

The accession number for the shotgun metagenomic data for the Irish COMBINE cohort reported in this paper is BioProject accession (NCBI): PRJEB77202. The accession number for the shotgun metagenomics data for the humanized-microbiota mice and re-sequencing of their donors as reported in this paper is BioProject accession (NCBI): PRJNA1413407.

The accession number for the shotgun metagenomic data for the Canadian CHILD cohort reported in this paper is BioProject accession (NCBI): PRJNA1403004. The informed consent obtained from the CHILD participants, in addition to the CHILD Inter-Institutional Agreement (IIA) which has been executed between the five Canadian institutions responsible for the study, govern the sharing of CHILD data. Data described in this manuscript are available by registration to the CHILD database (https://childstudy.ca/childdb/) and the submission of a formal request. All reasonable requests will be accommodated. More information about data access for the CHILD Cohort Study can be found at https://childstudy.ca/for-researchers/data-access/. Researchers interested in collaborating on a project and accessing CHILD Cohort Study data should contact.

Additional resources including bioinformatic workflows, analysis code, and supporting data are available at https://doi.org/10.5281/zenodo.17652611.

## ACKNOWLEDGEMENTS

We thank Mallory Ballard, Madeleine Castator, Patricia Gianeselo, Santosh Thapa, Funing Tian, Bansi Vanparia, Hector Acevedo, Chelsea Calderon, Margaret Conner, and Stephanie Fowler for assistance and support in the project. We are grateful for all mothers and infants who provided samples used in this study, and all staff who assisted with recruitment, enrollment, collection, and processing of samples. This work was supported by funding from the Wellcome Leap 1kD program to F.S.M, M.E.K, D.M.M, C.S., R.P.R., M.C.-M., K.F.T., R.A.B, and H.A.D. The COMBINE study was funded by a grant to M.E.K from Science Foundation Ireland (INFANT/B3067), which was co-funded by the European Regional Development Fund (ERDF) under Ireland’s European Structural and Investment Funds Programmes 2014–2020. This project was also supported in part by a grant from the Weston Family Foundation to C.P. and S.E.T., and by the Texas Medical Center Digestive Diseases Center (TMC-DDC; NIH P30DK056338) through services by the Cellular and Molecular Core, Functional Genomics and Microbiome (FGM) Core, and Gastrointestinal Experimental Model Systems Core which performed the gnotobiotic studies. T.D.H. is supported by an Office of Research Infrastructure Programs (ORIP)-based S10 Award (NIH S10OD036416), the TCMC-DDC FGM Core (NIH P30DK056338), and receives salary support from a P01 Program Project Grant (NIH P01AI152999). Data analysis was performed on the HPC cluster managed by the Biostatistics and Informatics Shared Resource (BISR) and supported by a National Cancer Institute award (NIH P30CA125123) and Institutional funds from the Dan L Duncan Comprehensive Cancer Center and Baylor College of Medicine.

## AUTHOR CONTRIBUTIONS

Conceptualization and study design: F.S.M., S.W.D., M.C.-M., K.F.T., R.A.B., and H.A.D.; Funding acquisition, F.S.M., C.S., R.P.R., M.E.K., D.M.M., C.P., S.E.T., M.C.-M., K.F.T., R.A.B., and H.A.D. Resources: C.P., S.E.T., C.S., R.P.R, M.E.K., and D.M.M. Identification and selection of infant samples: C.S., R.P.R., M.E.K., D.M. Sequencing of infant samples: G.J.A. Mouse husbandry, behavior assessment, dissection, and specimen collection: M.S., J.D.P., D.Z., C.K.A, A.B., N.M.R., J.S., and H.A.D. Brain dissection, immunohistochemistry, and imaging: D.-H.L., and Y.M. Targeted metabolomics methods and sample processing: T.D.H. Isolation of bacterial strains from infant feces: M.S. and E.C. Bioinformatic methods and analysis: F.S.M., A.K.A., and J.C. Metabolic modeling and analysis: F.S.M and H.A.D. Statistical analysis: F.S.M, D.-H.L., Y.M., J.C., R.J., D.L.Y.D., and H.A.D. Writing of original draft: F.S.M., K.F.T., R.A.B., and H.A.D. Review and editing of final manuscript: all authors.

## DECLARATION OF GENERATIVE AI AND AI-ASSISTED TECHNOLOGIES

Authors did not use any AI tool for the preparation or editing of this work and take full responsibility for the content of the publication.

## SUPPLEMENTAL INFORMATION

**Table S1. Tests for association between microbial species and pathways with BSID-III scores for infants in the COMBINE study.**

Microbial species and pathway abundances were tested for association with BSID-III scores using linear mixed effects models, including covariates of sequencing depth, infant age, infant sex, household income, household location (urban/rural), gestational age, delivery procedure (vaginal/Caesarean), birth weight, and feeding behavior (breastfed, formula-fed, or combination fed). To account for repeated sampling, individual subjects were included as a random effect. Each analysis was performed either on “All” samples (1-month-old to 12-months-old), “Early” samples (1-month-old to 4-month-old), or “Late” samples (9-month-old to 12-month-old) separately. “All” included 435 samples from 129 infants; “Early” included 234 samples from 125 infants; and “Late” included 117 samples from 84 infants.

**Table S2. Comparison of baseline clinical characteristics between low-scoring, typical-scoring, and high-scoring infants.**

**Table S3. Tests for association of microbial strains and pathways with BSID-III scores for a subset of infants in the COMBINE study, whose samples were used to establish humanized microbiota mouse lines.**

Here, we applied the same approach used for analysis in Table 1 to a smaller subset of samples. Each analysis was performed on “Early” or “Late” samples separately. Microbial strains correspond to the species genome-level bins (SGB) assigned by MetaPhlAn. “Early” included 57 samples for 31 infants and “Late” included 30 samples for 20 infants.

**Table S4. Tests for association of mouse microbial strains, pathways, neuroactive gene modules, and other features with donor BSID-III outcomes, related to Figure 4A**.

Microbial strains, pathways, KEGG orthologues, enzyme classes, and protein families were tested for association with BSID-III group (LE versus TE and HE) using linear mixed effects. Covariates included sequencing depth and infant age. To account for repeated sampling, mouse lines were included as a random effect. Microbial strains correspond to the species genome-level bins (SGB) assigned by MetaPhlAn. Microbial pathways and other features correspond to functional assignments by HUMAnN. Neuroactive gene modules were tested for association with BSID-III group (LE versus TE and HE) using feature set enrichment analysis, with ranks derived from the results of the linear mixed effects modeling of KEGG orthologs. Each analysis was performed on “Early” and “Late” samples separately. “Early” samples included 108 samples from 28 mouse lines and “Late” included 81 samples from 21 mouse lines.

**Table S5. Tests for association of mouse fecal metabolites and metabolic groups with donor BSID-III outcomes, related to Figures 4B-D.**

Abundance of individual metabolites, based on non-targeted metabolomics of mouse fecal pellets, were tested for association with BSID-III group (LE versus HE) using linear mixed effects models. Covariates included metabolomics batch and donor age. To account for repeated sampling, mouse lines were included as a random effect. Metabolic groups were tested for association with BSID-III group using feature set enrichment analysis, with ranks derived from the results of linear mixed effects modeling of individual metabolites and groups defined by meta-data provided by Metabolon. Each analysis was performed on “Early” and “Late” samples separately. “Early” included 54 samples from 18 mouse lines and “Late” included 39 samples from 13 mouse lines.

**Table S6. Tests for association of primary metabolic pathways and metabolite cross-feeding scores in mouse microbiomes with donor BSID-III outcomes, related to Figure S5E.**

Primary metabolic pathways, identified and quantified by gutSMASH and BiG-MAP, and metabolic exchange scores, estimated by MICOM, were tested for association with BSID-III group (LE versus TE and HE) using linear mixed effects models including donor age as a covariate. To account for repeated sampling, mouse lines were included as a random effect and infant age was included as a fixed effect. Analysis was performed on “Early” and “Late” samples separately. “Early” included 108 samples from 28 mouse lines and “Late” included 81 samples from 21 mouse lines.

## SUPPLEMENTAL FIGURES TITLES AND LEGENDS

**Supplemental Figure 1.**
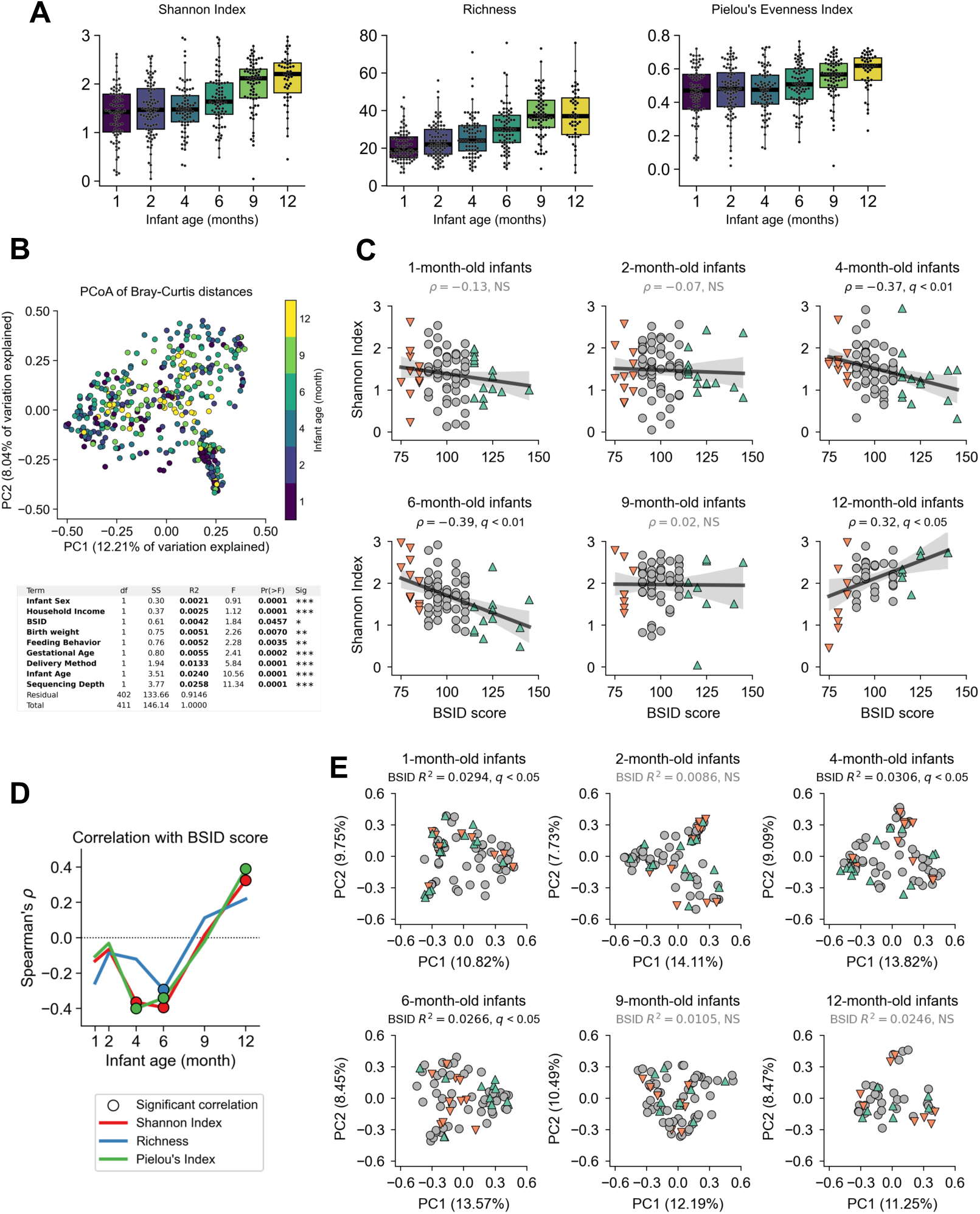
Microbial diversity in 129 infants of the COMBINE cohort. (A) Microbial alpha diversity metrics including Shannon index, Richness, and Pielou’s Evenness index, stratified by age. (B) Principal coordinates analysis of Bray-Curtis distance between infant samples with data points colored by infant age at time of sampling, and table summarizing result of PERMANOVA test on the Bray-Curtis distances using 9,999 permutations. (C) Correlation of Shannon index at different infant ages against infant cognitive score at 2 years old with a linear regression model fit. (D) Summary of correlation analyses for Shannon index (as shown in C), in addition to Richness and Pielou’s Evenness index. (E) Principal coordinates analysis of Bray-Curtis distance between infant samples at specific ages with data points colored by infant BSID-III cognitive score at 2 years old. Orange=low-scoring, gray=typical-scoring, green=high-scoring. Data set includes 426 samples for 129 infants. Data points stratified by age (month) were 1=75, 2=84, 4=75, 6=75, 9=71, 12=46 infants. Data are presented as medians with interquartile ranges (A). Statistical significance was determined by permutation testing (B and E) and Spearman correlation (C) and corrected with the Benjamini-Hochberg method. NS = not significant, q = adjusted p value.

**Supplemental Figure 2.**
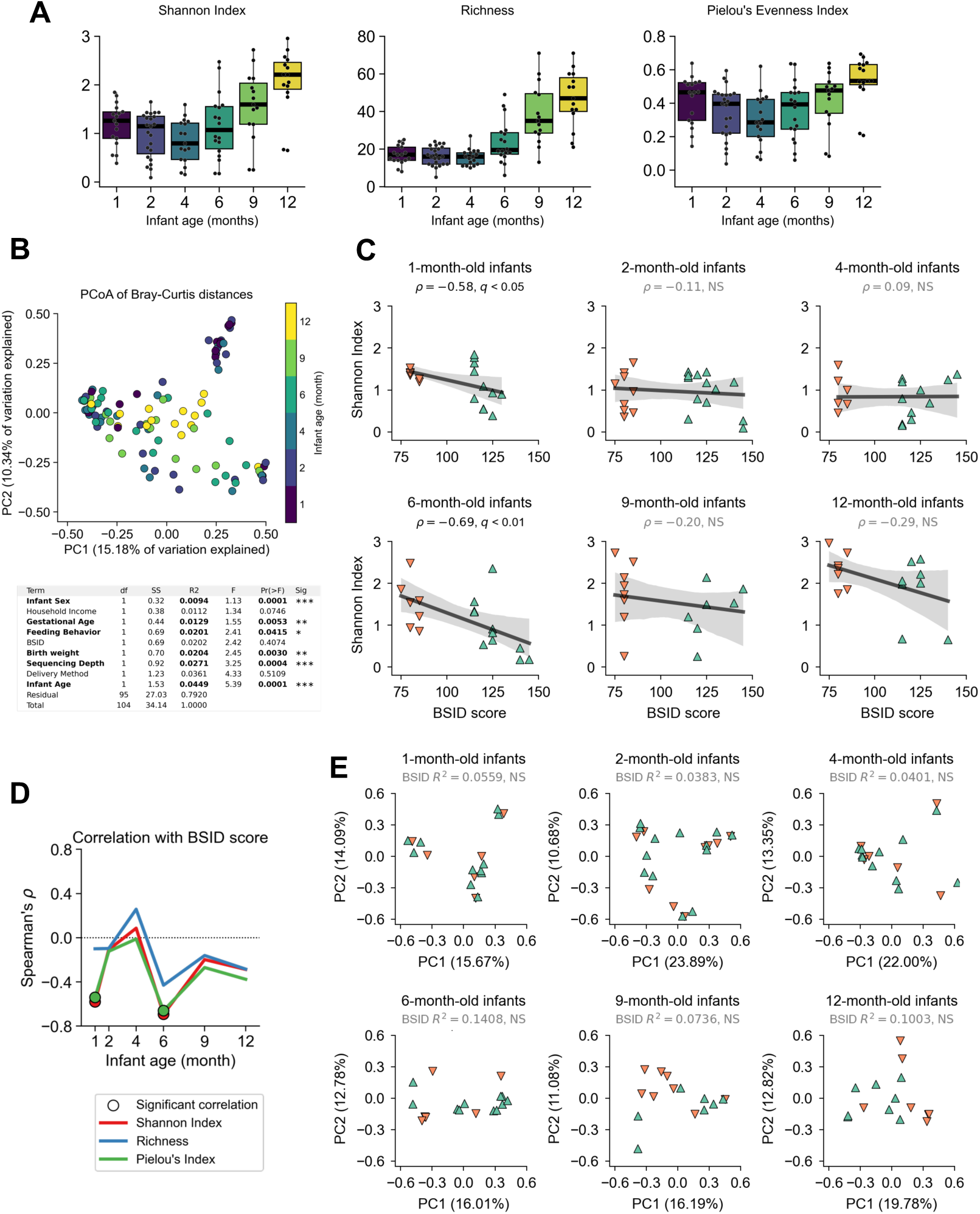
Microbial diversity in a subset of 32 infants of the COMBINE cohort whose samples established humanized-microbiota mouse colonies. (A) Microbial alpha diversity metrics including Shannon index, Richness, and Pielou’s Evenness index, stratified by age. (B) Principal coordinates analysis of Bray-Curtis distance between infant samples with data points colored by infant age at time of sampling, and table summarizing result of PERMANOVA test on the Bray-Curtis distances using 9,999 permutations. (C) Correlation of Shannon Index at different infant ages against infant cognitive score at 2 years old with a linear regression model fit. (D) Summary of correlation analyses for Shannon index (as shown in C), in addition to Richness and Pielou’s Evenness index. (E) Principal coordinates analysis of Bray-Curtis distance between infant samples at specific ages with data points colored by infant BSID-III cognitive score at 2 years old. Orange=low-scoring, gray=typical-scoring, green=high-scoring. Data set includes 105 samples for 32 infants. Sample sizes stratified by age (month) were 1=17, 2=23, 4=17, 6=18, 9=15, 12=15 infants. Data are presented as medians with interquartile ranges (A). Statistical significance was determined by permutation testing (B and E) and Spearman correlation (C) and corrected with the Benjamini-Hochberg method. NS = not significant, q = adjusted p value.

**Supplemental Figure 3.**
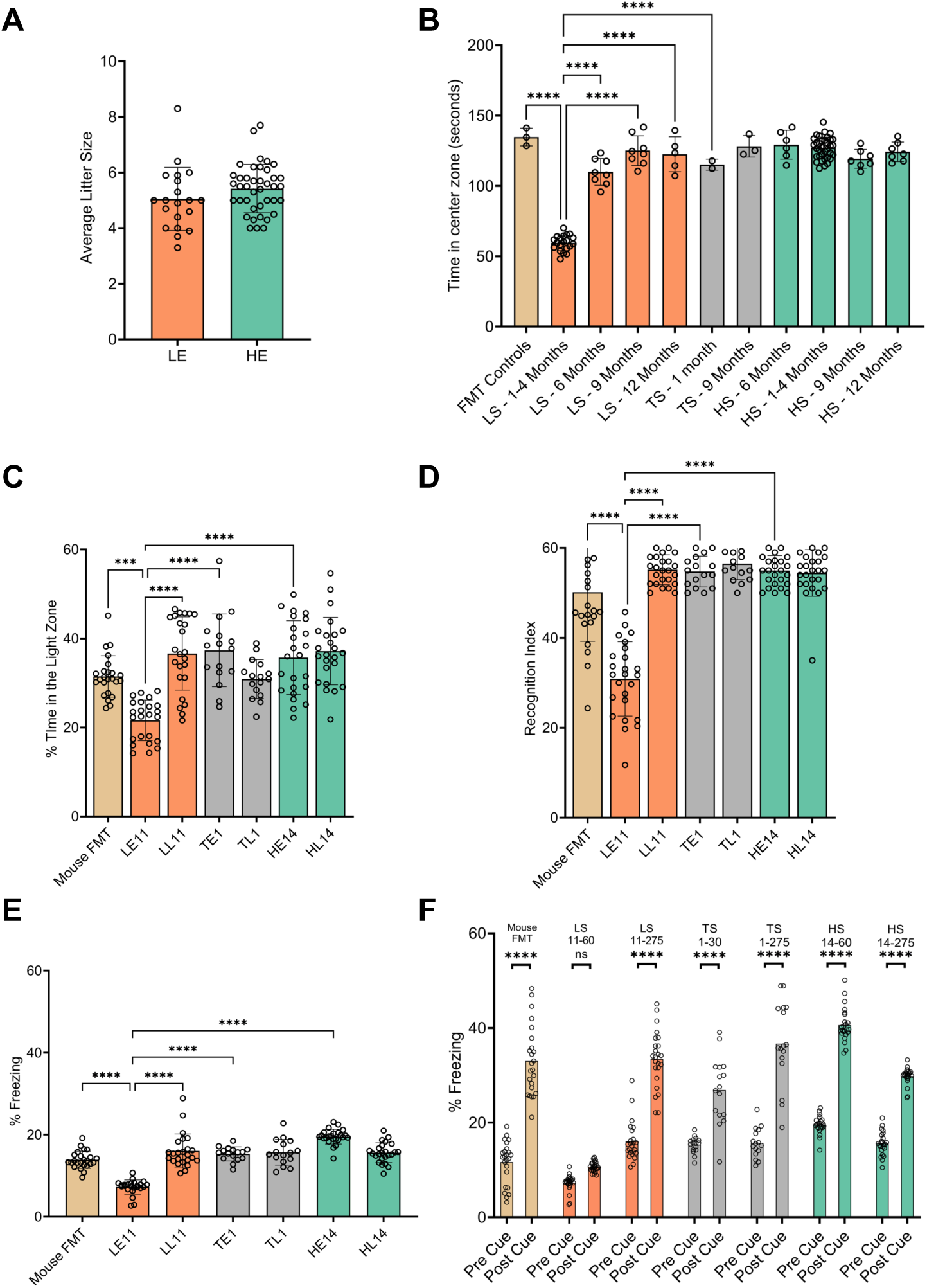
Additional mouse behavior results, related to Figure 2. (A) Average number of pups born to dams during breeding of litters 2–5. (B) Mice were given the open field behavioral test, for a total of 10 minutes, and the time spent in the center zone was quantitated. Each circle in the graph represents the average score of a mouse line (C) Mice were given the light dark box behavioral test, for a total of 10 minutes, and the time in either the light or dark compartment was quantitated. Each circle represents an individual mouse tested of a representative mouse line from indicated groups. (D) Mice were given the novel object recognition behavioral test, for a total of 10 minutes and the time spent with the novel object/total time was calculated and multiplied by 100 to report % of interaction time. Each circle represents an individual mouse tested for each representative mouse line of the indicated groups. (E-F) Box plot shows the percentage of total test time immobile during contextual (E) and cued (F) fear conditioning, which is the proxy for memory of adverse events following an auditory tone. Each circle represents an individual mouse tested for each representative mouse line of the indicated groups. Tan=controls, orange=low-scoring, gray=typical-scoring, green=high-scoring. In panel A, 1 point=24 mice tested, n=98 mouse lines. In panels B–F, 1 point=1 mouse, n=24. Data are presented as a mean ± SD. *p < 0.05, **p < 0.01, ***p < 0.001 **** p≤ 0.0001 by one-way ANOVA.

**Supplemental Figure 4.**
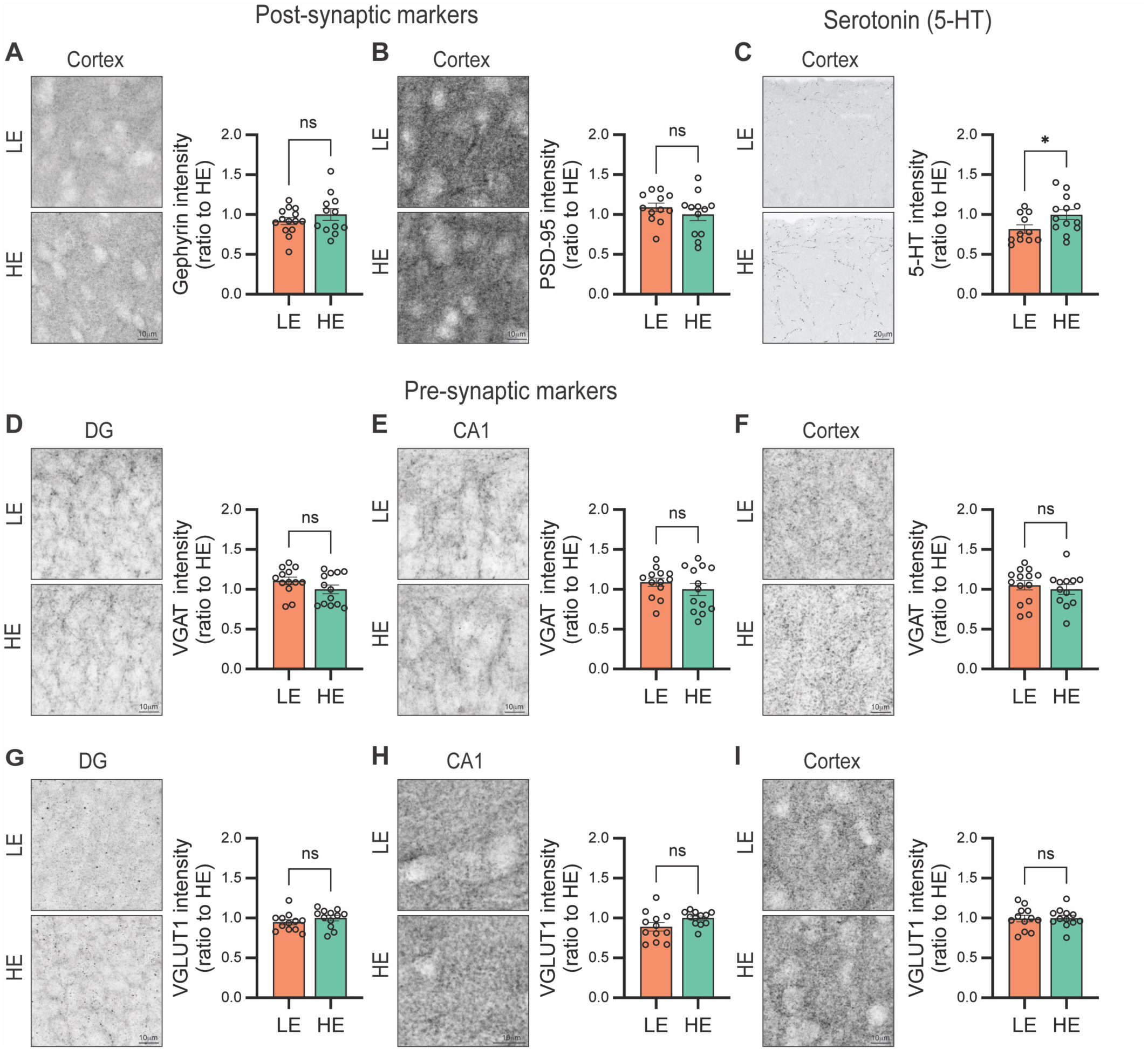
Additional brain data, related to Figure 3. (A–B) Postsynaptic inhibitory (Gephyrin) and excitatory (PSD95) marker expression in the somatosensory cortex showed no significant difference between LE and HE mice. (C) Serotonin (5-hydroxytryptamine, 5-HT) intensity was reduced in the somatosensory cortex of LE mice. (D–F) Inhibitory presynaptic development assessed by VGAT staining intensity in the dentate gyrus (DG), CA1, and somatosensory cortex showed no significant difference between LE and HE mice. (G–I) Excitatory presynaptic development evaluated by VGLUT1 intensity in the DG, CA1, and cortex showed no significant differences between LE and HE mice. Data are normalized to HE mice and presented as mean ± SEM. Ns, not significant, **p < 0.01, ***p < 0.001, ****p < 0.0001 by unpaired Student’s t-test.

**Supplemental Figure 5.**
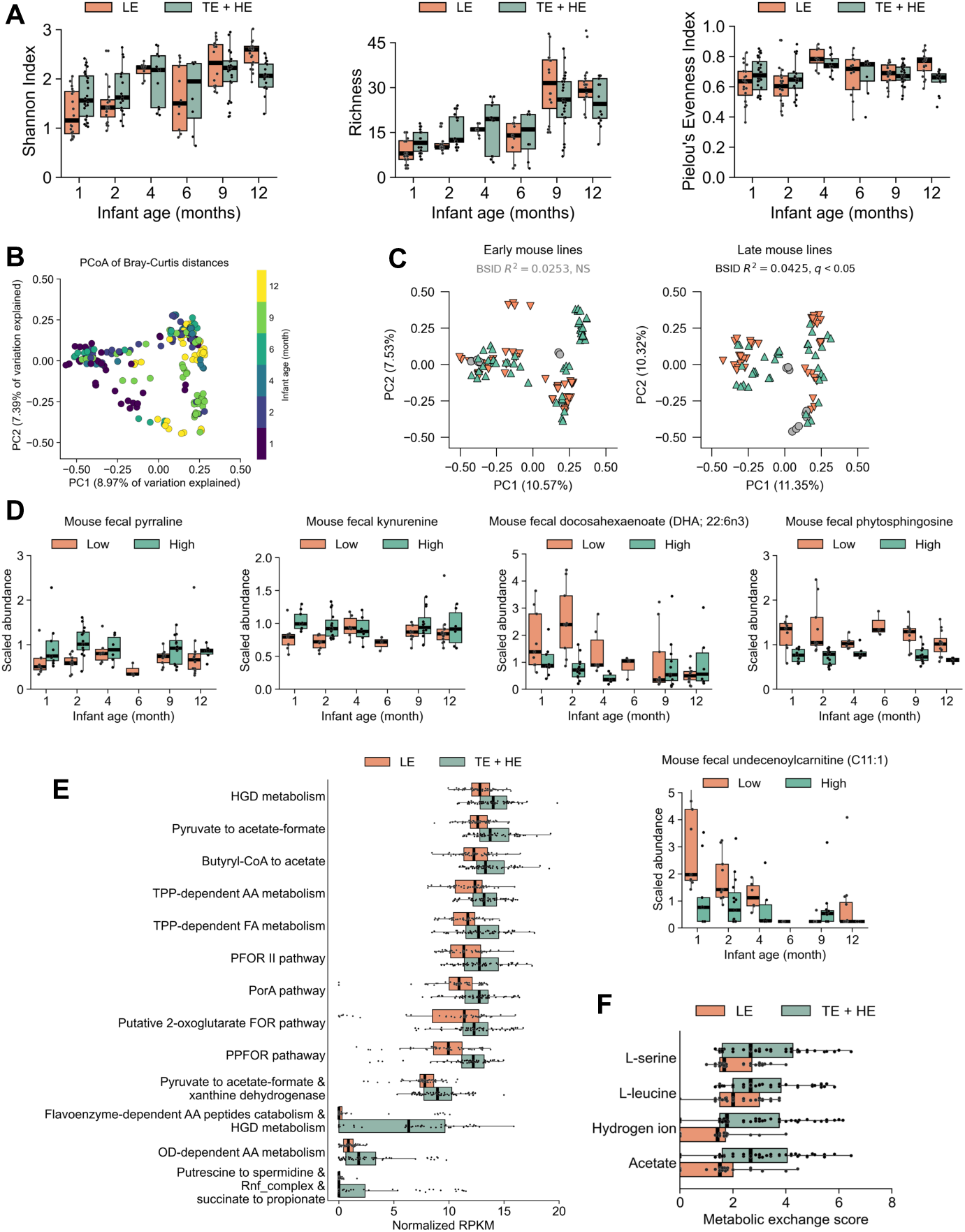
Microbial diversity and metabolic patterns in mice. (A) Alpha diversity metrics for mice stratified by donor’s age and BSID-III scores. There was no significant differences between groups at any age. (B) Principal coordinates analysis of Bray-Curtis distance between mouse samples with data points colored by infant age at time of sampling. (C) Principal coordinates analysis of Bray-Curtis distance between mouse samples humanized with either Early (1 to 4 months) or Late (9 to 12 months) infant samples. (D) Abundance of metabolites that are either significantly increased or decreased in LE compared to HE mice, related to Figures 4B-C and Table S5. (E) Relative abundance of primary metabolism pathways that were significantly depleted in LE mice, related to Table S6. (F) Metabolic exchange scores, or cross feeding scores, that were significantly depleted in LE mice in community-scale metabolic modeling, related to Table S6. AA=Amino acids, FA=Fatty acids, HGD=Homogentisate to 4-maleylacetoacetate, TPP=Thiamine pyrophosphate, PFOR=Pyruvate ferredoxin oxidoreductase, PPFOR=Phenylpyruvate ferredoxin oxidoreductase, OD=Oxidative decarboxylation. Orange=low-scoring, gray=typical-scoring, green=high-scoring. Data set is based on metagenomics for 209 mice with 3–4 mice per mouse line (A–C, E–F) and metabolomics for 96 mice with 3 mice per mouse line (D). Sample sizes stratified by age (month) are 1=48, 2=40, 4=20, 6=20, 9=45, 12=36 (A–C), 1=18, 2=24, 4=12, 6=3, 9=21, 12=18 infants (D), and 108 (E–F). For all panels, 1 point=1 mouse. Statistical significance was determined by Mann-Whitney U test (A), PERMANOVA (B and C), and linear mixed effects models (E–F) followed by Benjamini-Hochberg (A–F). In panels E-F, pathways or metabolites were visualized if they had nominal p < 0.05.

**Supplemental Figure 6.**
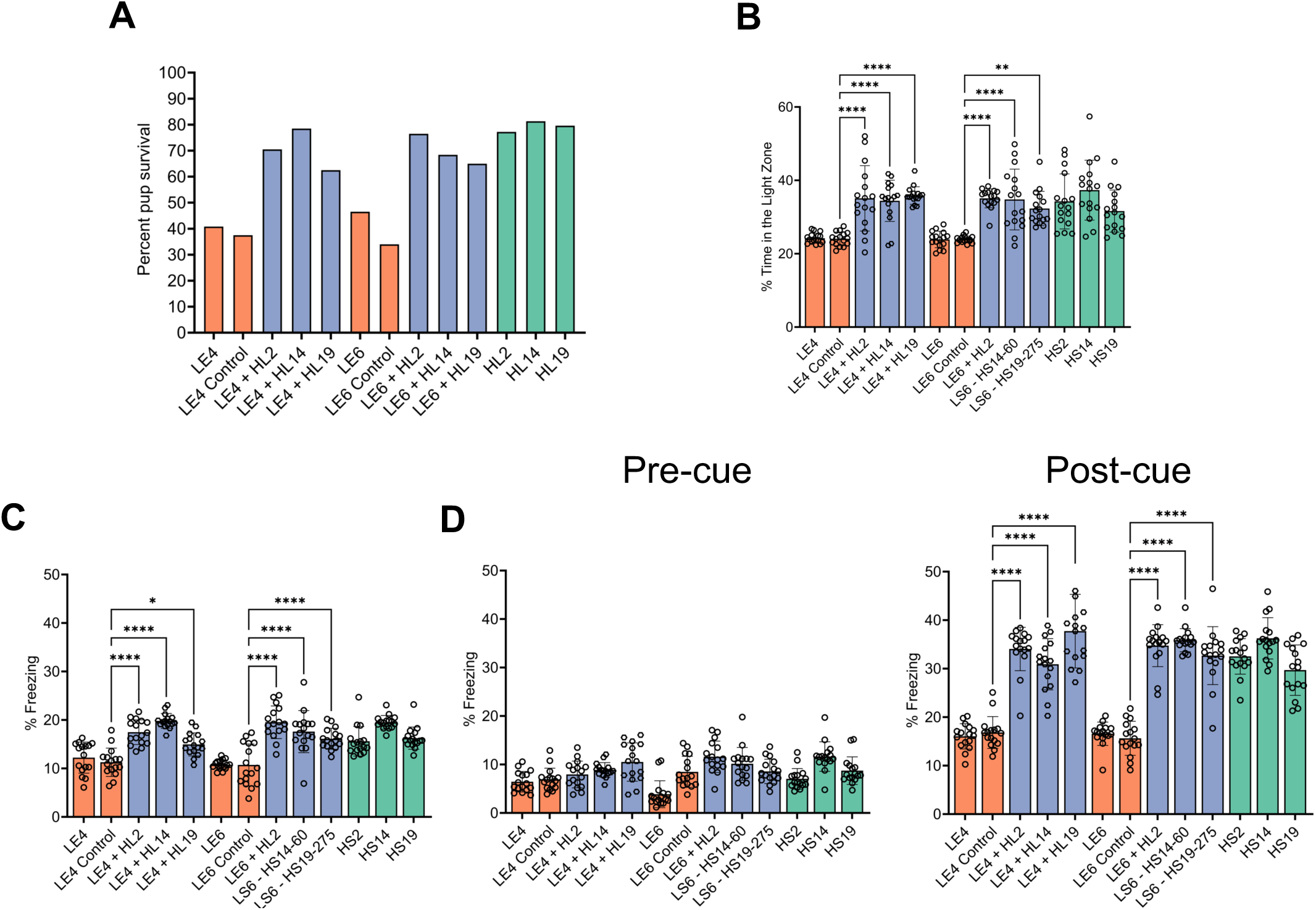
Additional FMT rescue data, related to Figure 5. (A) Pup survival returned to similar levels as HL animals when LE mice were given a second FMT from an HL infant at weaning. (B) Mice were given the light dark box behavioral test, for a total of 10 minutes, and the time in the light compartment was quantitated. Time exploring the light side increased relative to the LE controls. (C–D) Box plot showing the improvement in the contextual (D) and cued (E) fear conditioning test in LE animals that received a second FMT from a HL infant. Orange=low-scoring, blue=mice given a second FMT from HL infants, green=high-scoring. 1 point=1 mouse, n=16–24. Data are presented as a mean ± SD. *p < 0.05, **p < 0.01, ***p < 0.001, **** p≤ 0.0001 by one-way ANOVA.

**Supplemental Figure 7.**
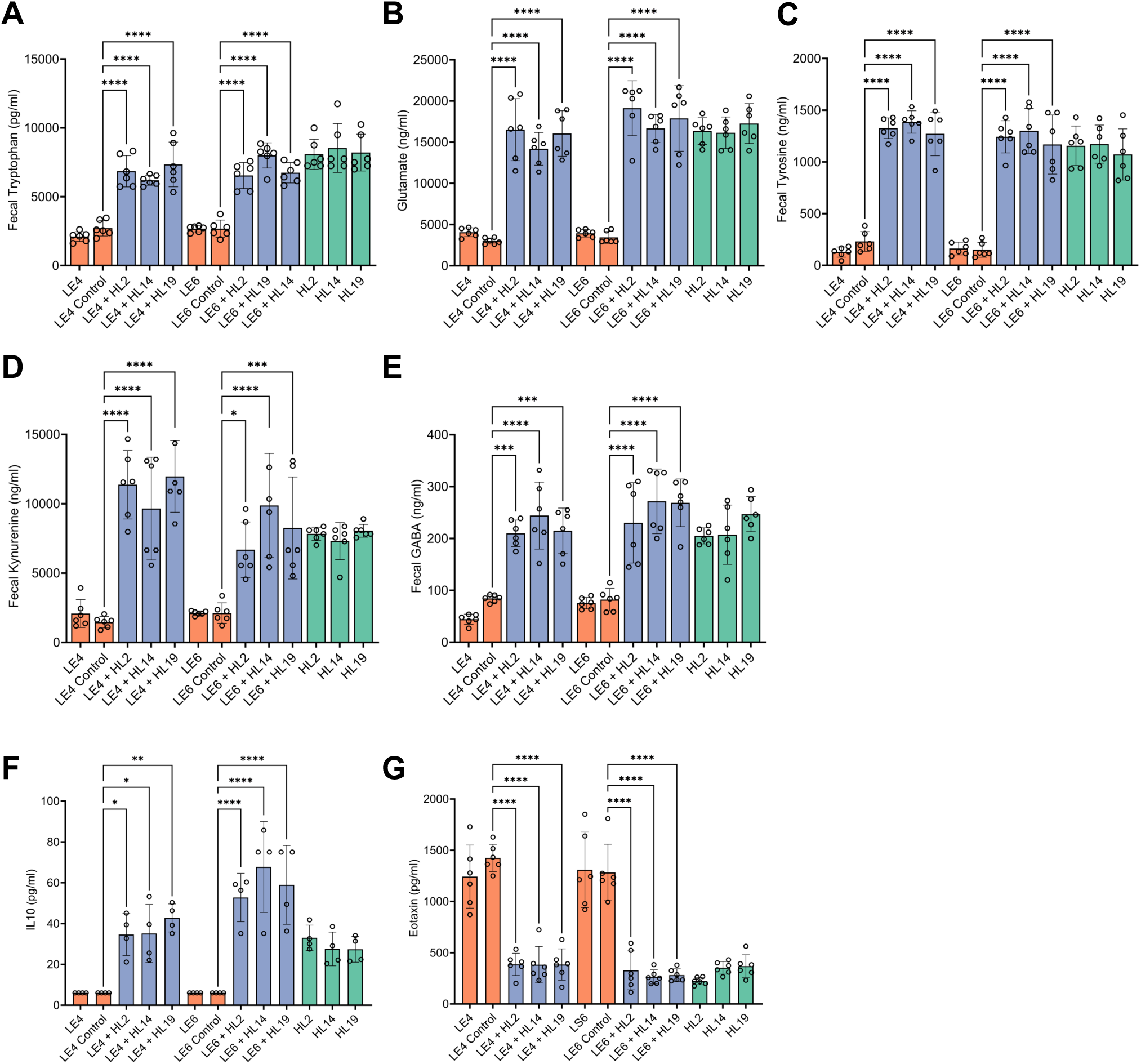
Additional FMT rescue data, related to Figure 5. (A–E) Fecal levels of tryptophan, glutamate, tyrosine, kynurenine, and GABA increased to comparable levels in HL animals. (F–G) Levels of IL-10 increased after second FMT to similar levels of HL mice. IL-10 was not detectable in LE mice (LOD of assay 6.06 pg/ml). Serum levels of proinflammatory eotaxin decreased in LE animals given a second FMT from HL infant. Orange=low-scoring, blue=mice given a second FMT from HL infants, green=high-scoring. In Panels A–E, 1 point=1 mouse, n=16–24. In panels F–G, 1 point=1 mouse, n=6 per group. Data are presented as a mean ± SD. *p < 0.05, **p < 0.01, ***p < 0.001, **** p≤ 0.0001 by one-way ANOVA.

**Supplemental Figure 8.**
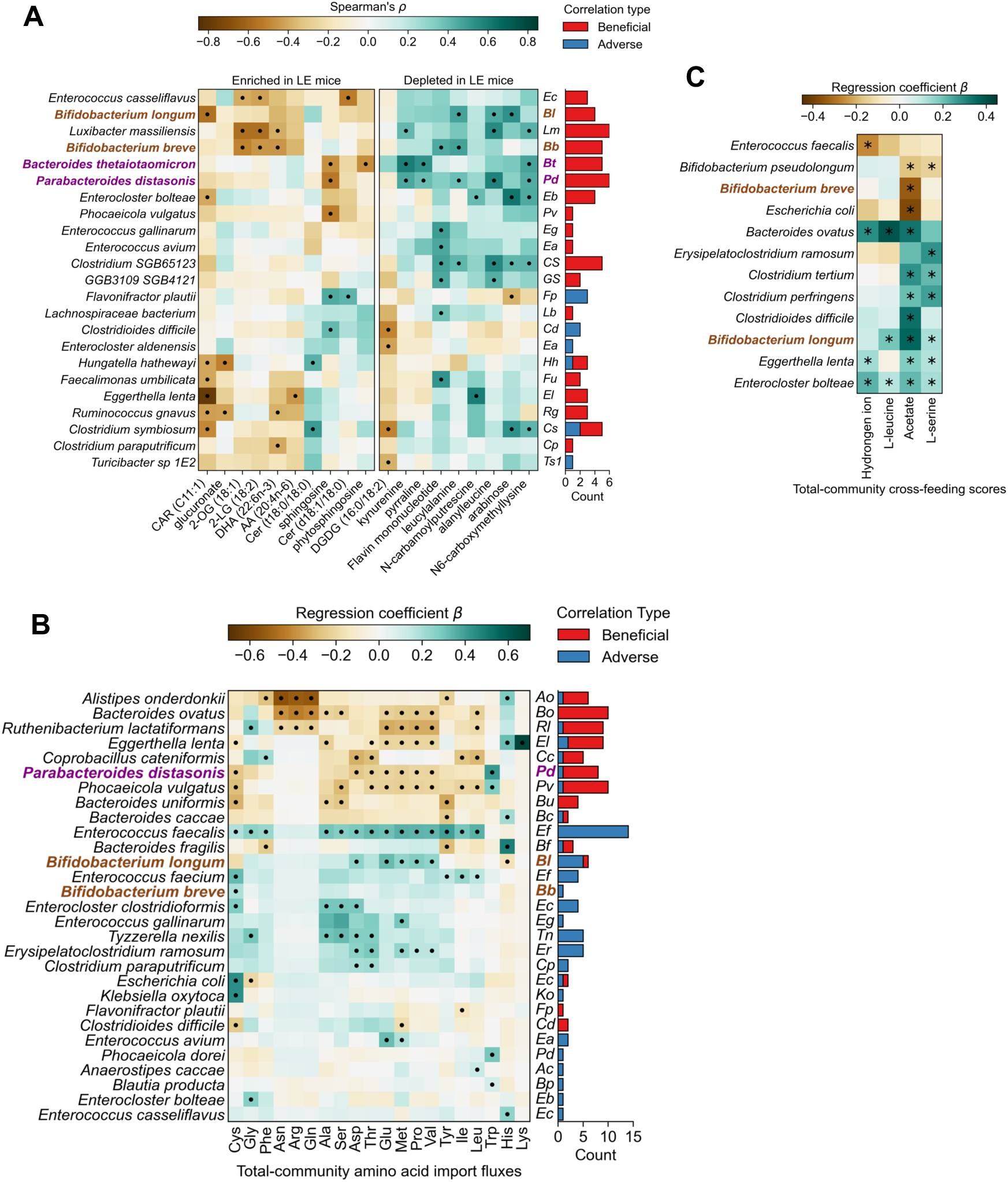
Correlation of microbes with metabolites and metabolic reactions in mice, related to Figure 6. (A) Correlations between microbial strains and metabolites that are enriched or depleted in LE mice. Correlations were tested using median abundances within each mouse. Metabolites were selected based on differential abundance analysis in Figure 4B and Table S5. (B) Correlations between species abundance and community-scale amino acid import fluxes (n=207 models). (C) Correlation between species abundance and metabolic exchange scores, which estimate cross-feeding of each metabolite. Metabolites included all differentially cross-fed metabolites between LE and HE mice as highlighted in Figure S4F and Table S6. Species labels in purple and brown indicate members of CS1 and CS2 respectively. Analysis was based on n=28 mouse colonies; n=3–4 mice per colony (A) and 207 metabolic models (B–C), and correlations were computed using Spearman’s test (A) or linear mixed effects models (B–C). • = nominal p < 0.05, * = adjusted p < 0.05 by Spearman correlation (A) or linear mixed effects models (B and C) and adjusted the Benjamini-Hochberg method (C).

## STAR★METHODS

### KEY RESOURCES TABLE

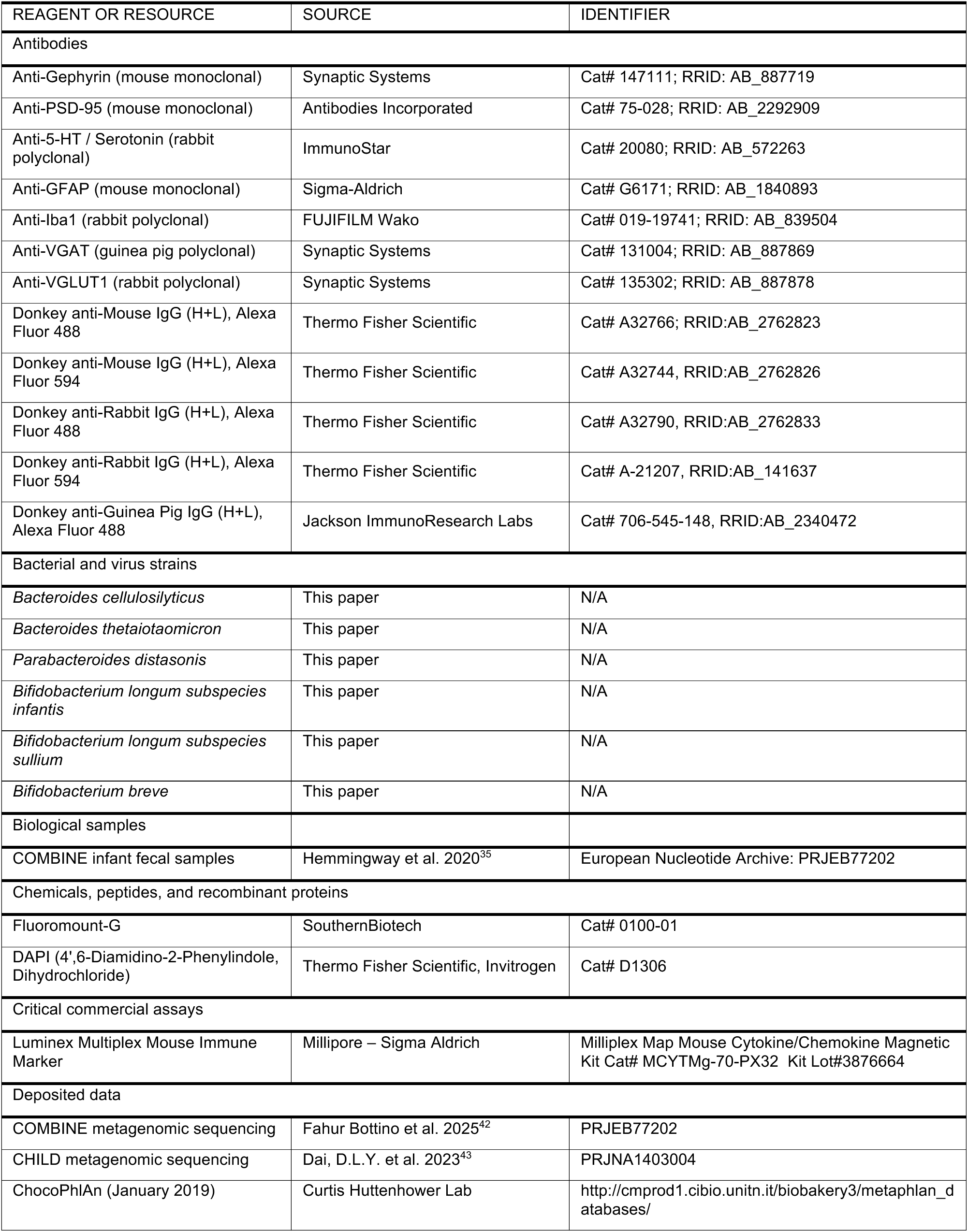
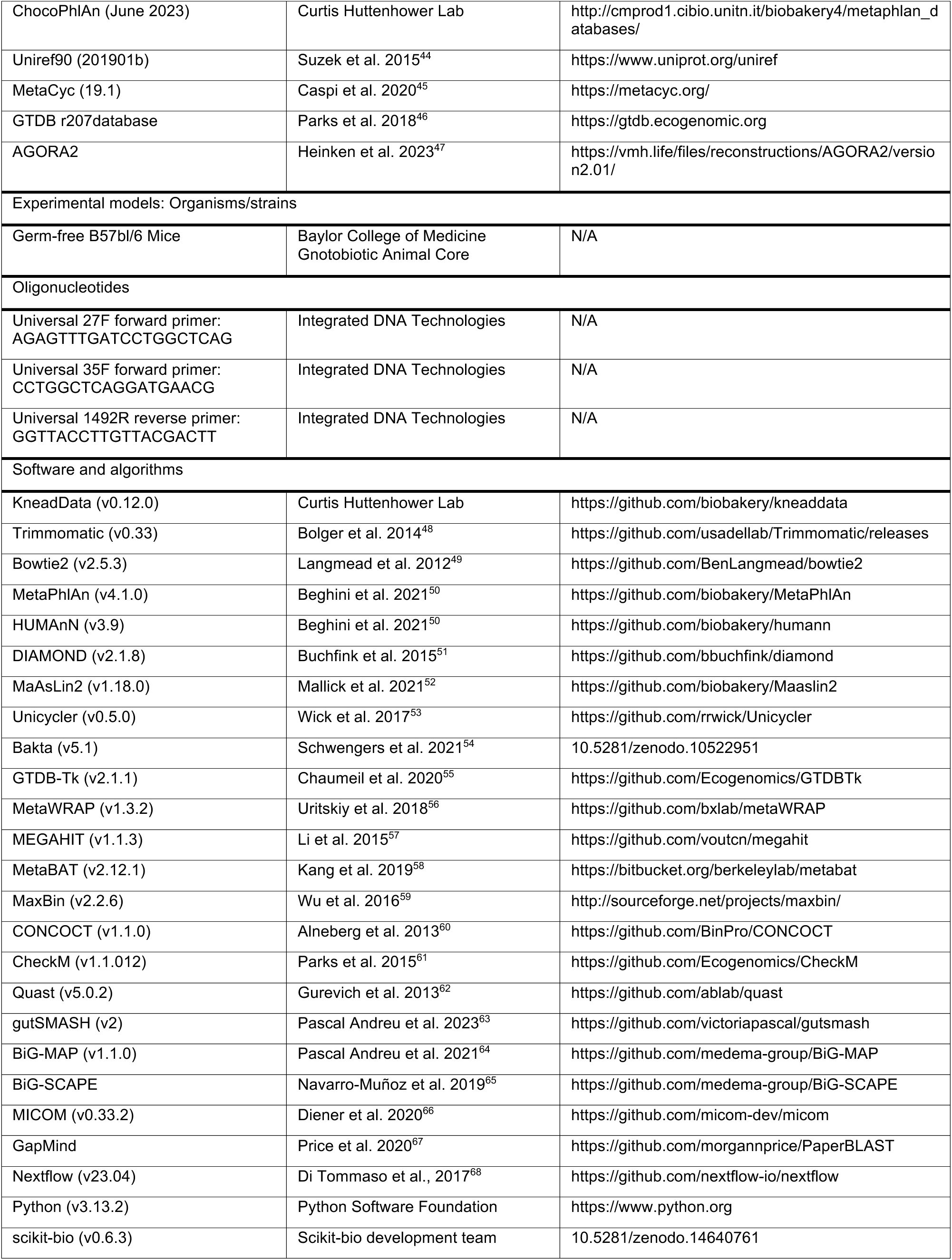
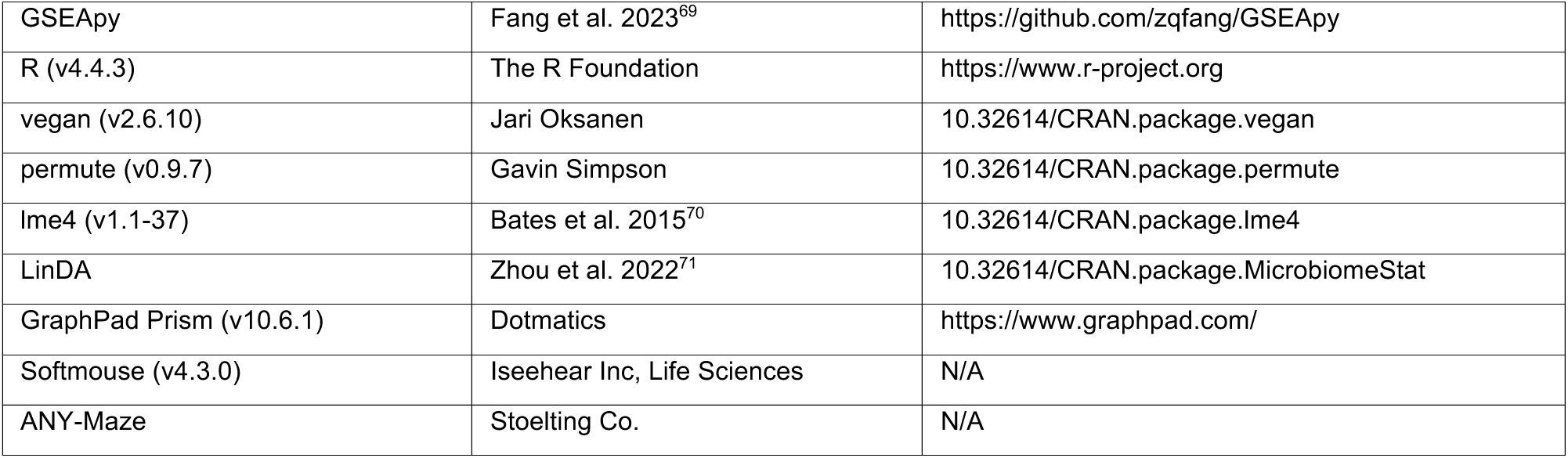

## EXPERIMENTAL MODEL AND STUDY PARTICIPANT DETAILS

### Description of the COMBINE cohort study

Infants in the COMBINE study were born to healthy females aged ≥16 years with a primiparous, low-risk, singleton pregnancy, who provided informed consent to participate in the IMPROvED Ireland pregnancy study and follow up COMBINE study. Ethical approval for COMBINE was granted by the Clinical Research Ethics Committee of the Cork Teaching Hospitals [ECM4(hh)06/01/15 and ECM3(bbb)10/04/18]. Infants were followed prospectively from birth through 2 years, at 1, 2, 4, 6, 9, 12, 18 and 24 months of age. Data on maternal demographics, pregnancy details, lifestyle, nutrition and health, delivery and early life events were obtained through interview and review of the medical records. Data on infant feeding, pediatric course, illness, medications, nutritional supplements, growth and developmental outcomes were collected at each postnatal study visit in the COMBINE study, as well as fecal samples, where possible. At 24 months, a psychologist conducted Bayley Scales of Infant and Toddler Development 3rd Edition (BSID-III), which is an assessment of developmental functioning consisting of 5 scales: cognition, receptive communication, expressive communication, fine motor, and gross motor. Published normative scores for the composite scales are a mean of 100 with an SD of 15; with developmental delay typically defined as a score <85.

### Whole-genome shotgun sequencing

Sequencing was performed at three different centers. Infant fecal samples from the COMBINE cohort were sequenced at the Teagasc Sequencing Center (Fermoy, Ireland). Resequencing of a subset of infant fecal samples was performed at the Baylor College of Medicine Alkek Center for Metagenomics and Microbiome Research (Houston, Texas). Sequencing of all mouse fecal pellets was performed at SeqCenter (Pittsburgh, Pennsylvania).

For all samples, DNA extraction was performed using the ZymoBIOMICS DNA Miniprep Kit. Sample libraries were prepared with the Illumina DNA Prep kit and sequenced on an Illumina NovaSeq or NextSeq platform, producing 2x151bp reads with 2Gbp per sample. Demultiplexing, quality control and adapter trimming was performed with bcl-convert.

### Microbial community analysis

Sequences were further processed with Kneaddata for additional quality trimming with Trimmomatic and removal of host reads with Bowtie2, by aligning reads against the human GRCh37or mouse C57BL/6NJ reference genomes. Microbial taxonomic composition was estimated with MetaPhlAn (v4.1.0) using the ChocoPhlAn (July 2023) database, while microbial functional composition was estimated with HUMAnN (v3.9). Reads were first mapped to the UniRef90 (January 2019) reference protein catalog using DIAMOND then gene families were assigned to microbial pathways using the MetaCyc (v19.1) pathways. Gene families were further grouped into enzymes, KEGG orthogroups, and protein families. Sequencing by Teagasc Sequencing Center was analyzed with the same tools but using earlier versions for MetaPhlAn (v3.1), HUMAnN (v3.8), and the ChocoPhlAn v31 (January 2019) database. Community diversity was analyzed with Python using the scikit-bio package. PERMANOVA was performed in R with the vegan package using 9,999 permutations and including the following covariates which are frequently associated with either BSID-III or gut microbiome: sequencing depth, infant age, infant sex, household income, gestational age, birth weight, delivery procedure, and breastfeeding history. In case of repeated sampling, permutations were created based on a block design using the permute package.

### Differential abundance analysis

For differential analysis of microbial composition, we used raw counts of species abundance as generated by MetaPhlAn and ran linear mixed effects models using LinDA, which imputed zeros based on library size, accounted for compositionality with center-log ratio transformation, and filtered species with prevalence of at least 10%. For microbial features quantified by HUMAnN, we performed differential abundance with linear mixed effects models on log-transformed abundances in R using MaAsLin2. For metabolic features, we performed differential abundance with linear mixed effects models in R using the lme4 package. For analysis of infant fecal samples, the model formula were “Microbial feature ∼ BSID Group + Infant Age + (1|Infant ID)”, with additional covariates that are frequently associated with either BSID-III or gut microbiome including infant sex, household income, gestational age, birth weight, delivery procedure, and breastfeeding history. For analysis of mouse samples, the model formula were “Microbial/metabolic feature ∼ Donor BSID Group + Donor Age + (1|Stool ID)”. Sequencing depth was added as a covariate for models analyzing microbial features and batch was added as a covariate for models analyzing metabolomic data. In each analysis, false discovery correction was performed on all p values for each fixed effect separately with the Benjamini-Hochberg method.

### Establishment of humanized microbiota mouse models

Gnotobiotic C57Bl6/J mice were obtained from the Center for Comparative Medicine at Baylor College of Medicine. Animals were kept a 12 hr light/dark cycle and had access to sterile food and water ad libitum. Infant fecal slurry was prepared by homogenization of frozen, solid infant feces in anoxic sterile PBS supplemented with 20% glycerol in an anaerobic chamber (90% N_2_, 5% CO_2_, 5% H_2_). A single breeding pair per fecal slurry was inoculated by oral gavage with a 100 µL volume of fecal slurry. They were maintained in a flexible isolator fed with HEPA-filtered air and provided with irradiated food and water until the dam was heavily pregnant, then animals were transferred to HEPA filtered Iso cages. Animals were then bred as independently stable mouse colonies with strictly sterile food, water, and handling prior to behavior experimentation. All behavioral tests were performed on 7- to 12-week-old mice, with particular care to avoid cross contamination between mouse lines by disinfecting surfaces and instruments with ammonium chloride wipes and 70% ethanol sanitization. Animal care and experimental procedures were approved by Baylor College of Medicine’s Institutional Animal Care and Use Committee in accordance with all guidelines set forth by the U.S. National Institutes of Health.

### Open field test

Open field testing was performed as previously described^72^. Mice were placed individually into a square Plexiglass arena (50 x 50 cm) with opaque walls and allowed to explore freely for 10 min under consistent dim lighting. Locomotor activity and anxiety-like behavior were quantified using AnyMaze tracking software, including total distance traveled and time spent in the center versus periphery. All testing was conducted during the light phase in a quiet, isolated room to minimize external stimuli.

### Light/dark box

Anxiety-like behavior was assessed using the light/dark box test as previously described^73^. Mice were placed in a two-chamber apparatus (40 × 40 cm) consisting of a brightly lit compartment and a dark, enclosed compartment connected by a small opening. Each mouse was introduced into the dark side and allowed to explore freely for 10 minutes. Time spent in each compartment and number of transitions were recorded using AnyMaze software. The apparatus was cleaned with 70% ethanol between trials to eliminate olfactory cues. Increased time in the light compartment was interpreted as reduced anxiety-like behavior.

### Novel object recognition

Recognition memory was assessed using the novel object recognition paradigm as previously described^74^. Mice were habituated to a 40 × 40 cm open field arena for 20 minutes. On day 2 (training), animals explored two identical objects placed in opposite corners of the arena for 10 minutes. On day 3 (testing), one familiar object was replaced with a novel object of similar size and complexity; object positions were counterbalanced across subjects to control for side bias. All objects were black; familiar objects were spheres and novel objects were cubes. Exploration was defined as nose-directed contact within a predefined interaction zone and was scored using AnyMaze software. A discrimination index was calculated to quantify preference for the novel object.

### Fear conditioning

Associative learning and memory were assessed using a two-day fear conditioning paradigm^75^. On day 1 (training), mice were placed in isolation chambers equipped with calibrated shock (0.7 mA) and sound (85 ± 2 dB) delivery systems. After a baseline period, a tone (conditioned stimulus) was paired with a mild foot shock (unconditioned stimulus) across three intervals (0–120 s, 150–210 s, 240–300 s). On day 2, contextual memory was tested by re-exposing mice to the original chamber without tone or shock, followed by a cued test in a novel chamber containing a vanilla-scented stimulus and altered bedding.

Freezing behavior was recorded and quantified using FreezeFrame software with thresholds set to 6 for training and 2 for cue testing. Apparatuses were cleaned thoroughly between microbiomes to prevent cross-contamination.

### Sociability and social novelty

Sociability and preference for social novelty were assessed using Crawley’s three-chamber test as previously described^14^. Mice were placed in a 60 × 40 × 23 cm Plexiglass arena divided into three interconnected chambers and allowed to habituate for 10 min. During the sociability phase, subjects explored freely and interacted with either an empty wire cup or a cup containing an age- and sex-matched unfamiliar conspecific (mouse 1); cup placement was counterbalanced across trials. In the social novelty phase, a second unfamiliar mouse (mouse 2) was introduced into the previously empty cup. Time spent in each chamber and direct interaction with each stimulus was recorded using AnyMaze software and scored by independent observers.

### Animal tissue processing

Mice were deeply anesthetized with Euthasol (200 mg/kg, Virbac) and transcardially perfused with ice-cold phosphate-buffered saline (PBS, pH 7.4), followed by 4% paraformaldehyde (PFA) in PBS. Brains were carefully removed and post-fixed in 4% PFA overnight at 4°C. Tissue was then cryoprotected in 30% sucrose in PBS for 48–72 h at 4 °C until fully equilibrated. Cryoprotected brains were embedded in Optimal Cutting Temperature (OCT; Tissue-Tek) compound in cryomolds and rapidly frozen. Coronal sections (40 µm) were cut on a cryostat (HM525NX, Thermo Scientific) and stored at –20 °C in cryoprotectant solution (50% glycerol / 50% PBS) until use.

### Immunohistochemistry

Free-floating coronal sections were washed in PBS and incubated in blocking solution (5% donkey serum and 0.3% Triton X-100 in PBS) for 1 h at room temperature (RT). Sections were then incubated with primary antibodies diluted in blocking solution overnight at RT. Following incubation, sections were washed three times for 10 min in PBS and incubated with fluorophore-conjugated secondary antibodies for 1 h at RT in the dark. The following antibodies were used: mouse anti-Gephyrin (Synaptic Systems, 147111; 1:500), mouse anti-PSD-95 (Antibodies Inc., 75-028; 1:500), rabbit anti-5-HT (ImmunoStar, 20080; 1:5000), mouse anti-GFAP (Sigma-Aldrich, G6171; 1:1000), rabbit anti-Iba1 (Wako, 019-19741; 1:1000), guinea pig anti-VGAT (Synaptic Systems, 131004; 1:1000), rabbit anti-VGLUT1 (Synaptic Systems, 135302; 1:1000), Alexa Fluor 488 donkey anti-mouse IgG (Thermo Fisher, A32766; 1:500), Alexa Fluor 594 donkey anti-mouse IgG (Thermo Fisher, A32744; 1:500), Alexa Fluor 488 donkey anti-rabbit IgG (Thermo Fisher, A32790; 1:500), Alexa Fluor 594 donkey anti-rabbit IgG (Thermo Fisher, A-21207; 1:500), and Alexa Fluor 488 donkey anti-Guinea Pig IgG (Jackson ImmunoResearch Labs, 706-545-148; 1:500). After three additional 10-min PBS washes, nuclei were counterstained with DAPI (Invitrogen), and sections were mounted on glass slides using Fluoromount-G (SouthernBiotech) and coverslipped.

### Image acquisition

Images were acquired using a Zeiss Apotome system equipped with 10× (NA 0.3) or 20× (NA 0.8) dry objectives, or a Zeiss LSM 880 confocal microscope equipped with 40× (NA 1.3) and 63× (NA 1.4) oil-immersion objectives, operated in Airyscan Fast mode. For synaptic marker imaging, Z-stacks were collected with a 0.5-µm optical step size to cover the 10-µm thickness of each section. Laser power, gain, digital offset, and detector settings were kept constant across all samples within each experiment. Images were acquired at 2048 × 2048 pixel resolution and saved in uncompressed .czi format for quantitative analysis.

### Image processing and quantification

Signal intensity quantification was performed in Fiji (ImageJ, NIH) using the original .czi files acquired from the microscope. All images within an experimental cohort were collected using identical acquisition parameters and processed using a uniform analysis workflow. Regions of interest (ROIs) corresponding to defined anatomical areas (e.g., cortex, hippocampus) were manually outlined using the Polygon Selection tool based on established landmarks. Mean fluorescence intensity for each ROI was obtained using Analyze > Measure. Background values were measured from a nearby non-specific region and subtracted from the corresponding ROI. Background-subtracted intensities were normalized to the highest value within each dataset to generate relative intensity units.

### Untargeted metabolomics

Untargeted metabolic profiling was performed by Metabolon (Durham, NC, USA). Mouse fecal pellets were freshly collected from mice, flash frozen, then stored at −80°C. Proteins were precipitated with methanol under vigorous shaking for 2 min (Glen Mills GenoGrinder 2000) followed by centrifugation. Resulting extract was divided into multiple fractions for analysis by two separate reverse phase (RP)/UPLC-MS/MS methods with positive ion mode electrospray ionization (ESI) and by HILIC/UPLC-MS/MS with negative ion mode ESI. Samples were placed briefly on a TurboVap (Zymark) to remove the organic solvent, then stored overnight under nitrogen before preparation for analysis. Because samples were split into multiple batches, a pooled matrix sample generated by taking a small volume of each experimental sample was included as a quality control (QC) sample spaced evenly among the injections across the platform run. Raw data was extracted, peak-identified and QC processed using a combination of Metabolon developed software services. Compounds were identified by comparison to library entries of purified standards or recurrent unknown entities. Biochemical identification was based on three criteria: retention index within a narrow retention time/index (RI) window, accurate mass to charge (m/z) ratio match to the library ± 10 ppm, and the MS/MS forward and reverse scores between the experimental data and authentic standards. Peaks were quantified using the area under the curve. Because samples were processed in multiple batches, block correction was applied by dividing raw peak areas by the median raw peak area for the QC samples in that batch, if metabolites are present in at least 50% of the QC samples. Zeroes for each metabolite were imputed with the metabolite’s lowest non-zero median-corrected value.

### Enrichment Analysis

We applied feature set enrichment analysis to detect groups of related neuroactive genes that are either enriched or depleted in LE mice compared to HE mice^37,76^. Sets with a minimum size of four genes were included in the analysis. Neuroactive genes were ranked based on a t-statistic and *p* values were computed using 999 permutations. We also performed metabolic group enrichment analysis by grouping metabolites into functionally relevant sets based on the “super pathway” and “sub pathway” annotations provided by Metabolon, the untargeted metabolomics vendor. Sets with a minimum size of four metabolites were included in the analysis. We ranked metabolites based on the differential abundance analysis between HE and LE mice (Table S5), by the t-value computed for each model using the lmerTest package. All enrichment analyses were performed with Python using the gseapy package and p values were computed using 999 permutations and corrected with the Benjamini-Hochberg method.

### Mouse plasma and fecal sample processing for targeted metabolomics

Under isoflurane anesthesia, blood samples were collected by cardiac puncture into EDTA, gently mixed by inversion, placed immediately on ice, and centrifuged at 3,000 RPM (4°C) for 15 minutes. A 20 µL volume of each individual mouse plasma sample was aspirated from the supernatant and placed into fresh 0.6 mL microcentrifuge tubes for the protein precipitation (PPT) procedure. An 80 µL volume of chilled methanol (−20°C) was added to each sample tube. Samples are vortexed for 30 seconds each to ensure complete PPT and then centrifuged at 17,000 *g* for 5 mins. The clarified serum extract supernatant was transferred into a fresh 0.6 mL microcentrifuge tube and stored frozen at −80°C until the samples could be transferred to the bioanalytical laboratory. After euthanasia, the large intestines are harvested and separated from the cecum. Feces are gently removed from the large intestine, weighed, and collected in 2 mL Fastprep tubes. Chilled methanol (−20°C) was added to yield a fecal density of 0.1 mg/µL. Tubes are homogenized by three cycles of bead beating for 1 minute followed by resting for 1 minute. After homogenization, tubes were centrifuged at 12,000 *g* for 5 minutes, and a volume of the clarified fecal extract supernatant was transferred into a fresh 0.6 mL microcentrifuge tube and stored frozen at −80°C until the samples could be transferred to the bioanalytical laboratory.

### SCIEX QTRAP 6500 and QTRAP 7500-Based LC-MS/MS Systems

A hybrid triple-quadrupole/linear ion trap mass spectrometer (QTRAP) 6500 (SCIEX, Framingham, MA, USA) was used to perform quantitative bioanalysis for the targeted tryptophan pathway and kynurenine-indole methods. The QTRAP 6500 MS system was connected to a Nexera Series-20 X2 UHPLC system (Shimadzu, Kyoto, Japan). System operation was performed using the Analyst® software (Version 1.6.2; SCIEX), while peak integration and quantitative analysis was performed using the MultiQuant™ software (Version 3.0.1; SCIEX).

A QTRAP 7500 was used to perform bioanalysis for the targeted proteogenic amino acid method and the combined arginine-polyamine pathway and glutamate cycle method. The QTRAP 7500 MS system was connected to a Shimadzu Series-40 Nexera UHPLC system. The system was operated using SCIEX OS software (Version 3.3.1.43; SCIEX), while peak integration and quantitative analysis was performed using the Analytics Module in SCIEX OS.

### Targeted tryptophan pathway method

The targeted tryptophan pathway method has been described previously^77,78^. Individual samples were prepared by mixing 10 µL volumes of clarified serum or stool extract supernatants into 90 µL volumes of an internal standard solution A (ISS-A) for a dilution factor (DF) of 10-fold that contained deuterated-metabolite internal standards (IS) concentrations of 500 ng/mL each for d5-tryptophan, d4-serotonin and d4-melatonin, and 1,500 ng/mL for d5-5-hydroxyindoleacetic acid (HIAA) prepared in water. Each sample was vortex-mixed for 30 seconds and transferred to an autosampler vial and a 10 µL volume of sample was injected onto the QTRAP 6500-based LC-MS/MS system. Instrument calibration for each metabolite was performed using calibration standards prepared at concentrations that spanned a linear dynamic range of 0.977-1,000 ng/mL for the following unlabeled metabolites: tryptophan, serotonin, melatonin, 5-HIAA, 5-hydroxytryptophan, N-acetylserotonin, tryptamine, and indoleacetic acid.

### Targeted kynurenine-indole method

The targeted kynurenine-indole method has been described previously^79^. Individual samples were prepared by mixing 10 µL volumes of clarified serum or stool extract supernatants into 90 µL volumes of an ISS-A (DF=10-fold) that contained deuterated-metabolite IS concentrations of 250 ng/mL each for d7-indole, d6-kynurenine, and d5-phenylalanine prepared in water. Each sample was vortex-mixed for 30 seconds and transferred to an autosampler vial and a 10 µL volume of sample was injected onto the QTRAP 6500-based LC-MS/MS system. Instrument calibration was performed using calibration standards prepared at concentrations 0.977-1,000 ng/mL for the following unlabeled metabolites: indole, kynurenine, phenylalanine, 3-hydroxykynurenine, and kynurenic acid.

### Targeted arginine-polyamine pathway and glutamate cycle method

The targeted, combined arginine-polyamine and glutamate Cycle method has been described previously^79^. Individual samples were prepared by mixing 10 µL volumes of clarified serum or stool extract supernatants into 190 µL volumes of an ISS-A (DF=20-fold) that contained deuterated-metabolite IS concentrations of 250 ng/mL each for d7-arginine, d7-citrulline, d7-ornithine, d4-putrescine, and d8-spermidine prepared in a mobile phase A (MPA) solution consisting of acetonitrile/100 mM ammonium formate (pH 4.0)/water (83/10/7, v/v/v). Each sample was vortex-mixed for 30 seconds and transferred to an autosampler vial and a 10 µL volume of sample was injected onto the QTRAP 7500-based LC-MS/MS system. Instrument calibration was performed using calibration standards prepared at concentrations that spanned a linear dynamic range of 1.95-2,000 ng/mL for the following unlabeled metabolites: arginine, agmatine, citrulline, N-carbamoylputrescine, ornithine, putrescine, spermidine, glutamate, glutamine, and GABA.

### Targeted proteogenic amino acid method

The targeted proteogenic amino acid method has been described previously^79^. Individual samples were prepared by mixing 10 µL volumes of clarified serum or stool extract supernatants into 190 µL volumes of an ISS-A (DF=20-fold) that contained deuterium and carbon-13 labeled metabolite IS concentrations of 750 nM for [13C6, 15N2]-L-cystine and 1,500 nM for all other uniformly [13C,15N]-labeled amino acid IS components prepared in a solution consisting of acetonitrile/water/formic acid (89.5/9.5/1, v/v/v). Each sample was vortex-mixed for 30 seconds and transferred to an autosampler vial then a 10 µL volume of sample was injected onto the QTRAP 7500-based LC-MS/MS system. Instrument calibration was performed using calibration standards prepared at concentrations that spanned a linear dynamic range of 2.44-2,500 nM for unlabeled cysteine, and 4.88-5,000 nM for the rest of the proteogenic amino acids.

### Microbial genome and metagenome analysis

Metagenome-associated genomes were assembled using MetaWRAP. MAGs were first assembled using Megahit then binned with MetaBAT, MaxBin, and CONCOCT. Bins were refined then reassembled by MetaWRAP with thresholds of 50% for completion and 5% for contamination. Assembly quality was analyzed with CheckM and QUAST. GTDB-Tk was used to identify MAG taxonomy using the GTDB r207database, then genome assemblies were annotated with Bakta.

Metabolic gene clusters were identified using gutSMASH and quantified with BiG-MAP. First, metabolic MAGs annotated with Bakta were analyzed with gutSMASH. Using all metabolic gene clusters detected across all samples, BiG-MAP next identified representative gene clusters with BiG-SCAPE then gene cluster families were counted in each metagenomic sample using Bowtie2. Raw read counts were converted to reads per kilobase per million, then normalized using cumulative sum scaling to account for sparsity. Metabolic gene clusters were then grouped into pathways. To identify pathways that differed between LE and HE mice, we performed linear mixed effects modeling using the following formula: “Pathway ∼ BSID Group + Infant Age + (1|Stool ID)” and corrected p values for multiple comparisons using the Benjamini-Hochberg method.

Amino acid biosynthetic potential was quantified using GapMind for all MAGs assembled from infant metagenomic samples. GapMind assigns scores to each amino acid biosynthetic pathway based on the lowest score for any of its steps (subpathways). We converted scores of High, Medium, and Low to values of 3, 2, and 1 respectively. For each species, its amino acid biosynthetic potentials were computed as the average across MAGs for all strains detected in our infant dataset.

### Species-metabolite correlation analysis

Species-metabolite correlation was performed as follows: Species abundance was summarized by their medians within each mouse colony (4 mice sequenced per line). Based on untargeted metabolomics, quality-control normalized metabolite abundances were likewise summarized by medians within each mouse line (3 mice per line). Spearman correlations were then computed for all unique pairs of species (with prevalence > 25%) and metabolites.

### Microbial community-scale metabolic models

Microbial community-scale metabolic modeling was performed with MICOM. Communities were simulated for growth with a cooperative tradeoff of 0.8 in the AGORA2 growth medium that recapitulates the Western diet. For each community, models were constructed only for species with existing models in the AGORA2 database, and species with relative abundance of at least 1%. If a model did not converge, the least abundant species was pruned, and model optimization was repeated, such that species were iteratively pruned by lowest abundance until a model converged. Exchange fluxes that were not estimated by a converged model were assigned a flux of zero. To identify metabolic exchange scores that differed between LE and HE mice, we performed linear mixed effects modeling using the following formula: “Score ∼ BSID Group + Infant Age + (1|Stool ID)”. To identify species that are correlated with metabolic exchange scores, we performed linear mixed effects modeling using the following formula: “reaction ∼ species + (1|Stool ID)”. P values were corrected for multiple comparisons using the Benjamini-Hochberg method.

### Isolation of bacterial strains from infant fecal slurries

Fecal slurry aliquots were thawed briefly and processed entirely under anaerobic conditions; all consumables and plates were deoxygenated in an anaerobic chamber for ≥24 h prior to use. Serial 10-fold dilutions were prepared in anoxic PBS. For each dilution, a 100 µL volume was spread onto agar plates (two plates per dilution, one per media type) using a sterile T-spreader; sterile PBS was plated as a negative control. Plates were allowed to absorb sample agar-side up, then inverted and incubated anaerobically at 37 °C.

Plates were inspected and colonies picked at days 1, 3, 5 and 7 post-plating; multiple colonies (>5) of distinct morphologies were preferentially sampled and smaller colonies targeted on later days. Fresh agar plates were deoxygenated overnight before streaking. Individual colonies were transferred with sterile loops and streaked onto fresh agar (isolated quadrants/sections per plate) and incubated anaerobically at 37 °C until colonies formed; plates were held at room temperature until downstream processing as needed.

### Bacterial strain identification

For 16S identification, single colonies were picked into 15 µL sterile water in PCR tubes inside an anaerobic chamber and subjected to colony PCR using appropriate polymerase and cycling parameters. Following 16S identification, strains of interest were further identified by whole-genome sequencing using 2x150bp Illumina sequencing on a NextSeq platform by SeqCoast (Portsmouth, NH) with 200Mbp per sample. Read demultiplexing and trimming was performed using DRAGEN (v4.2.7). Sequences were assembled into genomes with Unicycler then taxonomy was identified by GTDB-Tk using the GTDB r207 database.

### Cryopreservation

Following species identification, strains were inoculated into liquid medium and incubated anaerobically at 37 °C until confluent (typically 16-24h). For long-term storage, microbial culture was mixed 1:1 with sterile, deoxygenated 50% glycerol in PBS (resulting in 25% glycerol concentration) and aliquoted into cryovials for storage at −80 °C.

### Description of the CHILD cohort study

The CHILD study is a prospective longitudinal birth cohort study, which enrolled 3,405 subjects since pregnancy from 4 largely urban study centers across Canada (Vancouver, Edmonton, Winnipeg, and Toronto) from 2008 to 2012^80^. The study included 3,263 subjects eligible at birth, who were born at a minimum of 34 weeks of gestation and had no congenital abnormalities. CHILD Study followed children prospectively and collected detailed information on environmental exposures and clinical outcomes using a combination of questionnaires and in-person clinical assessments. This project is based on 365 participants from the Edmonton, Alberta site who had the Bayley Scales of Infant and Toddler Development, Third Edition (BSID-III) administered in early life. CHILD study research complies with all relevant ethical regulations and was written and approved by the University of British Columbia, University of Manitoba, University of Toronto, McMaster University, BC Children’s Hospital, The Hospital for Sick Children, and Simon Fraser University. The Research Ethics Board Number is H07-03120.

Infant fecal samples from the CHILD Study Cohort were sequenced by Diversigen (Minneapolis, MN, USA) and CosmosID (Germantown, MD, USA). DNA was extracted using the MO Bio PowerSoil Pro kit with bead beating in 0.1 mm glass bead plates. DNA quality was confirmed using Quant-iT PicoGreen. Sequencing libraries were prepared and run on an Illumina NextSeq platform using single-end 1×150 bp reads. Raw reads were processed with KneadData for quality control. This included removal of low-quality reads (Q-score <30), trimming of short reads (<50 bp), adapter trimming, and host-read decontamination using the human genome reference database. Only samples with at least 100,000 high-quality, non-host reads were retained for downstream analysis. Taxonomic (species and strain) and functional profiling was performed using MetaPhlAn (v4.1) for taxonomic assignment and HUMAnN (v3.8) for pathway profiling.

## QUANTIFICATION AND STATISTICAL ANALYSIS

For breeding, behavioral, targeted metabolomics, and immune data sets, data are presented as means ± the standard deviations. Comparisons between groups were made with the one- or two-way analysis of variance (ANOVA), using the Tukey’s Multiple comparisons test. GraphPad was used to generate graphs and statistics (GraphPad Software, Inc., La Jolla, CA). A p value of <0.05 was considered significant, and “n” indicates the number of mice or mouse lines as indicated of experiments performed.

For brain data, processed data were exported to GraphPad Prism for statistical analysis. Normalized values are presented as mean ± SEM. Statistical significance was assessed using unpaired Student’s t test for comparisons between two independent groups. Significance thresholds were defined as: n.s. *p* > 0.05, *p < 0.05, **p < 0.01, ***p < 0.001, and ****p < 0.0001.

