## Supplemental Table 2 for "Infant gut microbiomes contribute to metabolic states that impact brain function"

### Baseline characteristics for 296 infants in the COMBINE study

| Variable | Subcategory | Overall | High BSID | Typical BSID | Low BSID | P.value | Test |
|---|---|---|---|---|---|---|---|
| n | | 296 | 66 | 203 | 27 | | |
| Sex, n (%) | Female | 128 (43.2) | 30 (45.5) | 92 (45.3) | 6 (22.2) | 0.069 | Chi-squared |
| | Male | 168 (56.8) | 36 (54.5) | 111 (54.7) | 21 (77.8) | | |
| Household location, n (%) | Urban | 178 (60.3) | 42 (64.6) | 120 (59.1) | 16 (59.3) | 0.727 | Chi-squared |
| | Rural | 117 (39.7) | 23 (35.4) | 83 (40.9) | 11 (40.7) | | |
| **Household income**, n (%) | ≤ 42,000 | 56 (19.4) | 9 (13.6) | 38 (19.2) | 9 (37.5) | **0.032** | Chi-squared |
| | 43,000 – 84,000 | 154 (53.5) | 40 (60.6) | 108 (54.5) | 6 (25.0) | | |
| | ≥ 85,000 | 78 (27.1) | 17 (25.8) | 52 (26.3) | 9 (37.5) | | |
| Gestational age, median [Q1,Q3] | | 40.4 [39.4,41.0] | 40.6 [39.7,41.1] | 40.4 [39.3,41.0] | 40.1 [39.1,40.9] | 0.17 | Kruskal-Wallis |
| Delivery method, n (%) | Vaginal | 214 (72.8) | 52 (78.8) | 147 (73.1) | 15 (55.6) | 0.072 | Chi-squared |
| | C-section | 80 (27.2) | 14 (21.2) | 54 (26.9) | 12 (44.4) | | |
| Birth weight, mean (SD) | | 3520.1 (475.4) | 3610.6 (449.1) | 3500.6 (486.1) | 3445.6 (441.2) | 0.183 | One-way ANOVA |
| Feeding behavior, n (%) | Breastfed | 150 (60.0) | 36 (64.3) | 102 (60.0) | 12 (50.0) | 0.68 | Chi-squared |
| | Combination fed | 56 (22.4) | 13 (23.2) | 36 (21.2) | 7 (29.2) | | |
| | Infant formula fed | 44 (17.6) | 7 (12.5) | 32 (18.8) | 5 (20.8) | | |

### Baseline characteristics for a subset of 32 infants in the COMBINE study, whose samples established humanized-microbiota mouse lines

| Variable | Subcategory | Overall | High BSID | Low BSID | P.value | Test |
|---|---|---|---|---|---|---|
| n | | 32 | 20 | 12 | | |
| Sex, n (%) | Female | 8 (25.0) | 7 (35.0) | 1 (8.3) | 0.204 | Fisher's exact |
| | Male | 24 (75.0) | 13 (65.0) | 11 (91.7) | | |
| Household location, n (%) | Urban | 19 (59.4) | 13 (65.0) | 6 (50.0) | 0.473 | Fisher's exact |
| | Rural | 13 (40.6) | 7 (35.0) | 6 (50.0) | | |
| Household income, n (%) | ≤ 42,000 | 6 (18.8) | 2 (10.0) | 4 (33.3) | 0.102 | Chi-squared with Yate's correction |
| | 43,000 – 84,000 | 12 (37.5) | 10 (50.0) | 2 (16.7) | | |
| | ≥ 85,000 | 14 (43.8) | 8 (40.0) | 6 (50.0) | | |
| Gestational age, median [Q1,Q3] | | 40.6 [39.8,41.3] | 41.0 [40.1,41.4] | 40.1 [39.2,40.4] | 0.054 | Kruskal-Wallis |
| **Delivery method**, n (%) | Vaginal | 23 (71.9) | 17 (85.0) | 6 (50.0) | **0.049** | Fisher's exact |
| | C-section | 9 (28.1) | 3 (15.0) | 6 (50.0) | | |
| Birth weight, mean (SD) | | 3615.0 (505.1) | 3656.0 (492.7) | 3546.7 (539.8) | 0.573 | Welch's t-test |
| **Feeding behavior**, n (%) | Breastfed | 22 (68.8) | 17 (85.0) | 5 (41.7) | **0.017** | Chi-squared with Yate's correction |
| | Combination fed | 7 (21.9) | 3 (15.0) | 4 (33.3) | | |
| | Infant formula fed | 3 (9.4) | 0 (0.0) | 3 (25.0) | | |
